# Disruptions in Outer Membrane-Peptidoglycan Interactions Enhance Bile Salt Resistance in O-antigen-Producing *E. coli*

**DOI:** 10.1101/2025.02.09.637341

**Authors:** Jilong Qin, Yaoqin Hong, Waldemar Vollmer, Renato Morona, Makrina Totsika

**Affiliations:** Centre for Immunology and Infection Control, School of Biomedical Sciences, Queensland University of Technology, Brisbane, QLD, Australia; Max Planck Queensland Centre, Queensland University of Technology, QLD, Australia; Institute for Molecular Bioscience, The University of Queensland, Brisbane, QLD, Australia; Biomedical Sciences and Molecular Biology, College of Medicine and Dentistry, James Cook University, QLD, Australia; Centre for Bacterial Cell Biology, Biosciences Institute, Newcastle University, Newcastle upon Tyne, UK; School of Biological Sciences, Department of Molecular & Biomedical Sciences, Research Centre for Infectious Diseases, University of Adelaide, SA, Australia

**Keywords:** polysaccharides, cell envelope, O antigen, peptidoglycan, Lpp, OmpA

## Abstract

Bile salts (BS) are derived from cholesterol in the liver and act as antimicrobial agents in the intestines by disrupting bacterial cell membranes and inducing oxidative stress. The gut bacterium *E. coli* is naturally resistant to BS, including the model strain K12 that produces a truncated LPS without O-antigen (OAg). Paradoxically, restoring a wild-type like LPS with OAg sensitises *E. coli* K12 to exogenous BS. In this study, we investigate this phenomenon. We show that mutations causing truncation of the LPS core oligosaccharide render these strains even more susceptible to BS, similar to the mutant strain MG1655-SΔ*waaL* defective in OAg ligase, primarily due to the accumulation of the lipid-linked intermediate UndPP-OAg. Through the characterisation of BS-resistant suppressor mutants of MG1655-SΔ*waaL*, we identify key genetic disruptions involved in resistance. Notably, we observed the highest BS resistance in strains with a weaker connection between the outer membrane (OM) and peptidoglycan (PG), including strains lacking the major OM-anchored, PG-binding proteins OmpA or Lpp, or expressing versions of these that lack PG-binding. Our data suggest that BS-induced stress in OAg-producing *E. coli* is due to the spatial constraints between OM and PG, and that mutations disrupting OM-PG interactions alleviate this stress, enhancing BS resistance. These findings provide new insights into a major challenge *E. coli* faces in the gut environment where it needs to produce OAg for stable colonisation and resists BS. *E. coli* can only survive BS exposure by fine-tuning the connectivity between its cell envelope layers, which highlights a potential target for modulating bacterial responses to BS in the gut.

**Author summary:** Enteric bacteria residing in the human gut must withstand the host-derived antimicrobial agents, bile salts (BS), but the underlying resistance mechanisms are not fully elucidated. This study investigates the BS resistance mechanisms in O-antigen (OAg)-producing *Escherichia coli* K-12. We show that truncation of lipopolysaccharide (LPS) core oligosaccharides or restoration of OAg production increases BS sensitivity due to the accumulation of UndPP-OAg intermediates. By analysing suppressor mutants, we identify key genetic disruptions, particularly affecting the level of outer membrane-peptidoglycan (OM-PG) interactions involving OmpA and Lpp, which confer heightened BS resistance. Our findings highlight how BS-induced stress is linked to spatial constraints between the OM and PG layer, offering new insights into bacterial adaptation to BS stress. This research may provide new targets for therapeutic interventions to modulate gut microbial responses to BS.

## Introduction

Bile salts (BS), cholate and chenodeoxycholate are derived from cholesterol in the liver and secreted from gallbladder storage into the intestines in humans. There, the two primary BS are modified by anaerobic bacteria through dehydroxylation to generate the secondary BS deoxycholate (DOC) and lithocholate, respectively [1]. BS are amphipathic molecules that act as detergents to help emulsifying fats, but they also possess potent antimicrobial activity by altering the permeability of the bacterial cell membrane [2, 3], unfolding cellular proteins [4] and inducing oxidative damage to DNA [5]. However, enteric bacteria have adapted to the human gut and resist the antimicrobial activities of BS, a key characteristic that was exploited in the development of the selective media MacConkey agar for the isolation and identification of gut bacteria [6]. Some bacterial pathogens including adherent-invasive *Escherichia coli* [7], *Shigella flexneri* [8] and *Salmonella enterica* [9] even evolved to utilise BS as an environmental clue to modulate their virulence function.

The major BS resistance mechanism in the model enterobacterium *E. coli* is through the exclusion of intracellular accumulation via multidrug efflux system such as AcrAB-TolC and MdtM [10]. One might expect to be able to identify direct cellular targets of BS from an *E. coli* mutant lacking the major efflux pump TolC. In one study [11], mutations conferring BS resistance in *E. coli* K-12Δ*tolC* were mapped to the *pstI-cyaA-crp* region, which encodes a (PTS)-cAMP-Crp regulatory cascade. These mutations reduced the carbohydrate metabolism and thus the accumulation of reactive oxygen species (ROS) [12], which can damage macromolecules like DNA [13], indicating that BS-induces a widely shared, oxidative stress-mediated death pathway. BS was also reported to upregulate chaperones in *Enterococcus faecalis* [14], suggesting that it induces a protein folding stress. Indeed, BS was shown to effectively cause protein aggregation and induce disulfide stress in *E. coli* K-12 lacking the cytosolic chaperone Hsp33 [4]. However, it remains unclear whether these consequences arise from a direct attack on key proteins or are downstream effects resulting from the interaction of BS with unidentified target(s).

As a model enterobacterium, *E. coli* K-12 has been used frequently in studying BS resistance. However, due to a mutation in the *wbbL* gene, K-12 strains do not produce OAg [15], a virulence determinant that is present in most newly isolated *E. coli* and important to colonise the human gut [16]. We have recently shown that the restoration of OAg production in *E. coli* K-12 sensitises it to the large antibiotic vancomycin in the presence BS [3], an effect that was also reproduced in other OAg-producing uropathogenic *E. coli* and *S. flexneri* strains, suggesting that BS induced cell envelope stresses when OAg is produced. Prolonged exposure of the OAg-restored *E. coli* K-12 selected for mutations leading to the inactivation of OAg production [3], confirming that the production of OAg poses the major stress in the presence of BS. The BS sensitisation effect in the OAg-producing *E. coli* was found to be due to the accumulation of the lipid-linked OAg intermediate, undecaprenol pyrophosphate-OAg (UndPP-OAg), since a Δ*waaL* mutant was found to be sensitive to BS on its own when OAg is being produced [3]. It was proposed that the accumulated UndPP-OAg intermediate caused disruptions in the biosynthesis of the cell envelope, leading to bacterial cells that are more susceptible to BS. However, the exact cell envelope processes affected remain to be elucidated.

Here, we show that mutations leading to the truncation of LPS core oligosaccharide in OAg-producing *E. coli* K-12 are all sensitive to BS, which can be attributed to the accumulation of UndPP-OAg in these strains. To understand the underling mechanism of increased BS sensitivity in these *E. coli* K-12 strains accumulating OAg intermediates, we selected for and characterised suppressor mutants capable of growing in the presence of a lethal dose of BS. Suppressor mutations conferring the highest BS resistance disrupted OAg biogenesis, and the non-OAg disrupting mutations were mapped primarily in genes responsible for outer membrane (OM) biogenesis. Importantly, mutations disrupting the peptidoglycan (PG)-interacting domain of OmpA or Lpp, which both reduce the linkage between OM and PG, conferred the highest resistance to BS. Our data suggest that BS poses stress to *E. coli* K-12 accumulating polymerised UndPP-OAg intermediates through periplasmic spatial constraint, and therefore the reduction in of OM-PG connections confers higher level of BS resistance.

## Results

### Accumulation of UndPP-OAg in periplasm sensitises E. coli to BS

The bacterial OM acts as a permeability barrier primarily enabled by LPS molecules on the outer leaflet, which limits the access of many antimicrobials to their intracellular targets [17]. Although LPS is essential, mutants with altered or truncated core structures remain viable. We therefore first investigated BS resistance of *E. coli* K-12 MG1655 with defects in LPS core oligosaccharide through deletion mutations abolishing inner core (Δ*waaC,* Δ*waaF* and Δ*waaP*) and outer core (Δ*waaB,* Δ*waaO,* Δ*waaG and* Δ*galU*) biosynthesis and assembly (Fig 1A), which were all confirmed to have truncated LPS core oligosaccharide (S1A Fig). Consistent with a previous report [18], only the heptose-less mutants Δ*waaF* and Δ*waaC* resulted in reduced BS resistance, and disruptions to outer core oligosaccharide and phosphorylation on heptose I (HepI) had no effect on BS resistance (Fig 1B).

**Fig 1.**
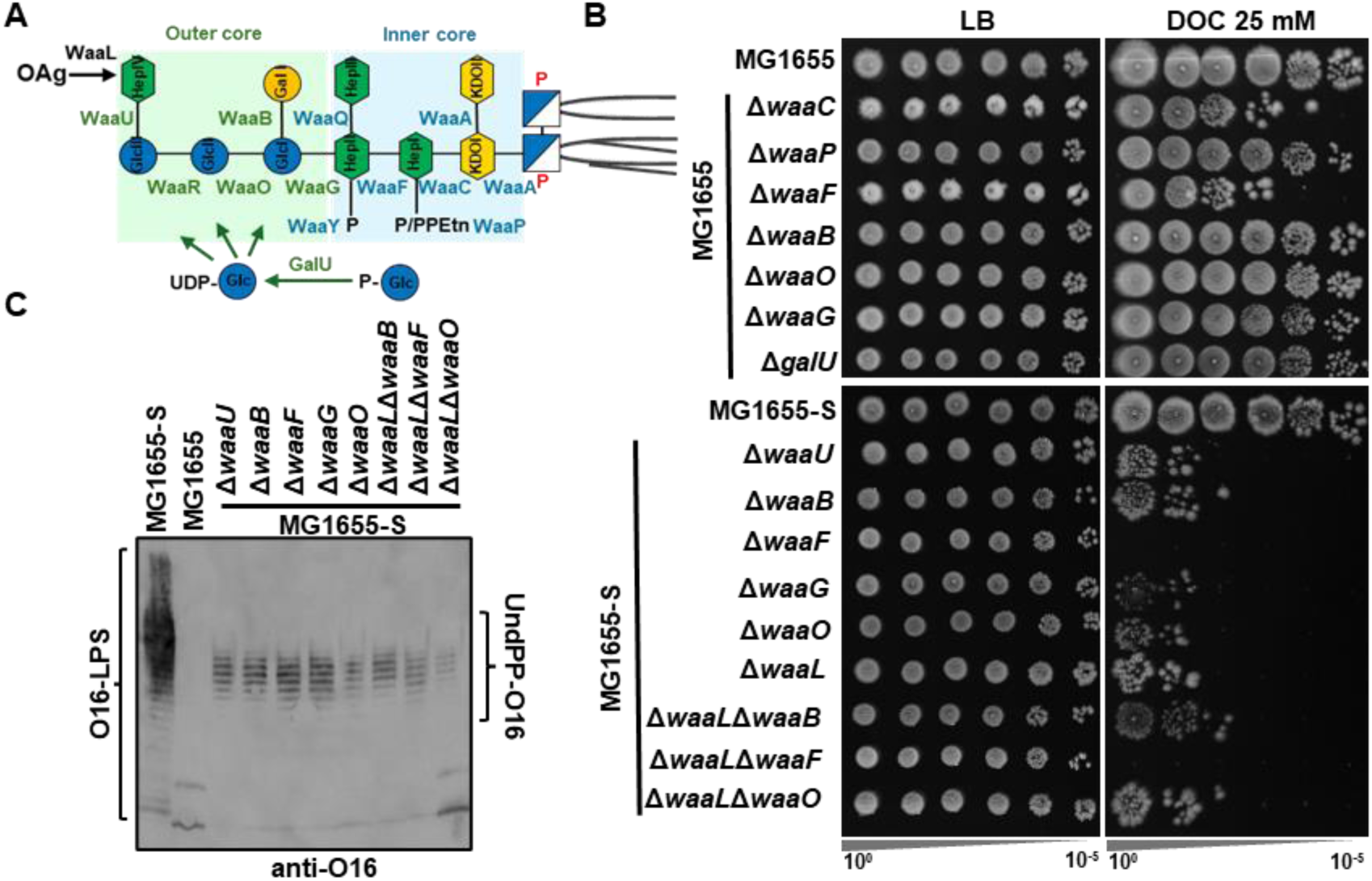
Accumulation of UndPP-OAg sensitises *E. coli* K-12 MG1655 to BS. **(A)** Schematic representation of *E. coli* K-12 LPS core oligosaccharide structure and assembly with enzymes for its assembly shown. **(B)** BS sensitivity assay for indicated *E. coli* K-12 strains. Overnight bacterial culture was adjusted to OD_600_ of 1 and spotted (4 µl) in 10-fold serial dilutions (10^0^-10^-5^) onto LB agar supplemented with or without 25 mM sodium deoxycholate (DOC). **(C)** Western immunoblotting of O16 OAg with lysate prepared from whole bacterial cells of indicated *E. coli* K-12 strains. Accumulated cellular UndPP-OAg are shown and labelled.

We have previously removed the *IS5* insertional element from *wbbL* in strain MG1655 to construct the OAg-producing *E. coli* K-12 strain MG1655-S [3] (S1B Fig). Disruptions of both LPS inner core (Δ*waaF*) and outer core (Δ*waaB,* Δ*waaO,* Δ*waaG and* Δ*waaU*) in MG1655-S resulted in truncation of LPS core oligosaccharide and loss of OAg capping due to the loss of distal HepIV required for OAg addition (S1B Fig). All tested outer core oligosaccharide assembly mutants in OAg-producing MG1655-S showed a drastic reduction in BS resistance (Fig 1B), while in OAg-deficient MG1655 this sensitivity was only observed in Δ*waaF* and Δ*waaC* mutants, suggesting that production of OAg sensitises mutants with LPS outer core truncation to BS. We have shown previously that an OAg ligase mutant Δ*waaL* had reduced BS resistance in MG1655-S due to accumulation of UndPP-OAg intermediates in the periplasm, but not in MG1655 [3]. Here, we show that all MG1655-S core truncation mutants accumulated UndPP-OAg (Fig 1C) and the level of reduction in BS resistance in outer core truncation mutants (Δ*waaB,* Δ*waaO,* Δ*waaG* and Δ*waaU*) was similar to that of Δ*waaL* mutant in the MG1655-S background (Fig 1B). Moreover, disruption of outer core (Δ*waaB,* and Δ*waaO*) in MG1655-SΔ*waaL* mutant had no further sensitisation to BS in comparison to corresponding outer core mutants in MG1655-S (Fig 1B), suggesting the sensitisation to BS in these MG1655-S outer core truncation mutants was primarily due to accumulation of UndPP-OAg intermediates.

The major known BS resistance mechanism is through exclusion of cellular BS via efflux pumps. However, it seems unlikely that the efflux pump activity was impaired when UndPP-OAg intermediates are accumulated in the periplasm, as MG1655-SΔ*waaL* was not further sensitised to the efflux pump substrate ampicillin [19] in comparison to MG1655 and MG1655-S (S1C Fig).

Interestingly, introducing inner core truncation through Δ*waaF* mutation in both MG1655-S and MG1655-SΔ*waaL* accumulated UndPP-OAg, further inhibited bacterial growth (over a 100-fold reduction than their respective parental strains) on the BS containing media (Fig 1B). This observation suggested that the reduction in BS resistance was due to the accumulation of UndPP-OAg intermediates and the defect in LPS heptose addition are additive and possibly through affecting different cellular components/processes. Disruption of the LPS inner core is known to severely compromise the formation of LPS-Ca^2+^ bridges which may weaken OM barrier function [20]. Counterintuitively, addition of Ca^2+^ instead slightly sensitised MG1655-SΔ*waaL* to BS, and chelation of Ca^2+^ by EGTA slightly restored its BS resistance (S1D Fig). These results suggest that the membrane barrier function empowered by LPS-Ca^2+^ bridges plays a limited role in the BS resistance and, in contrast, a reduced membrane stiffness may be beneficial to BS resistance when OAg is produced.

### Disruptions in OAg biosynthesis strongly enhances BS resistance in MG1655-SΔwaaL

To investigate the underlining mechanism of BS sensitisation effect due to the production and accumulation of UndPP-OAg intermediates, we harvested suppressor mutants (157 isolates) of MG1655-SΔ*waaL* that grew on LB agar media containing 2.5 mM DOC (DOC-LBA), designated as BP suppressor library (Fig 2A). All suppressor mutants grew on plain LB media with no observable growth defects (S1 Table). We then characterised growth kinetics of the BP library strains in liquid DOC-LB and ranked the BS resistance according to their culture optical density at 3.5 h (Fig 2A). A higher culture density was observed for all suppressor mutants in comparison to their parent strain MG1655-SΔ*waaL* (S1 Table). Disruption of OAg repeating unit (RU) biosynthesis and assembly restored BS resistance in MG1655-SΔ*waaL* [3]. We therefore first performed library screening to distinguish and exclude suppressor mutations affecting OAg biosynthesis and assembly by restoring the OAg-LPS (Smooth-LPS, S-LPS) production in the whole BP suppressor library via complementation of MG1655-SΔ*waaL*, resulting in a pWaaL-complemented BP suppressor library, designated as SBP sub-library (Fig 2A). S-LPS production protects *E. coli* MG1655-S from colicin E2 (ColE2) DNA endonuclease cell entry and bacteriophage P1 infection. We therefore patched SBP sub-library onto agar LB media containing ColE2 (ColE2-LBA) and P1kc phage (P1-LBA) as well as DOC-LBA (Fig 2A). All complemented SBP mutants retained BS resistance, growing on DOC-LBA with no observable defects, yet 14/157 SBP suppressor mutants showed growth defects on ColE2-LBA and 23/157 SBP suppressor mutants showed growth defects on P1-LBA (S1 Table). Consistent with our prediction, most SBP mutants with completely inhibited growth on ColE2-LBA and/or P1-LBA corresponded to BP mutants with the highest BS resistance amongst all mutants. Specifically, 11 SBP mutants exhibiting defects in ColE2-LBA and/or P1-LBA, corresponded to BP mutants ranked among the top 20 for BS resistance. (S1 Table). These 11 SBP mutants were further confirmed to lack detectable or altered S-LPS production (S2A Fig) and showed compromised resistance to ColE2 (S2B Fig). These results showed that our screening design was sufficiently robust and accurate to successfully exclude suppressor mutations affecting OAg RU assembly, and that the selected mutants retain BS resistance despite producing OAg.

**Fig 2.**
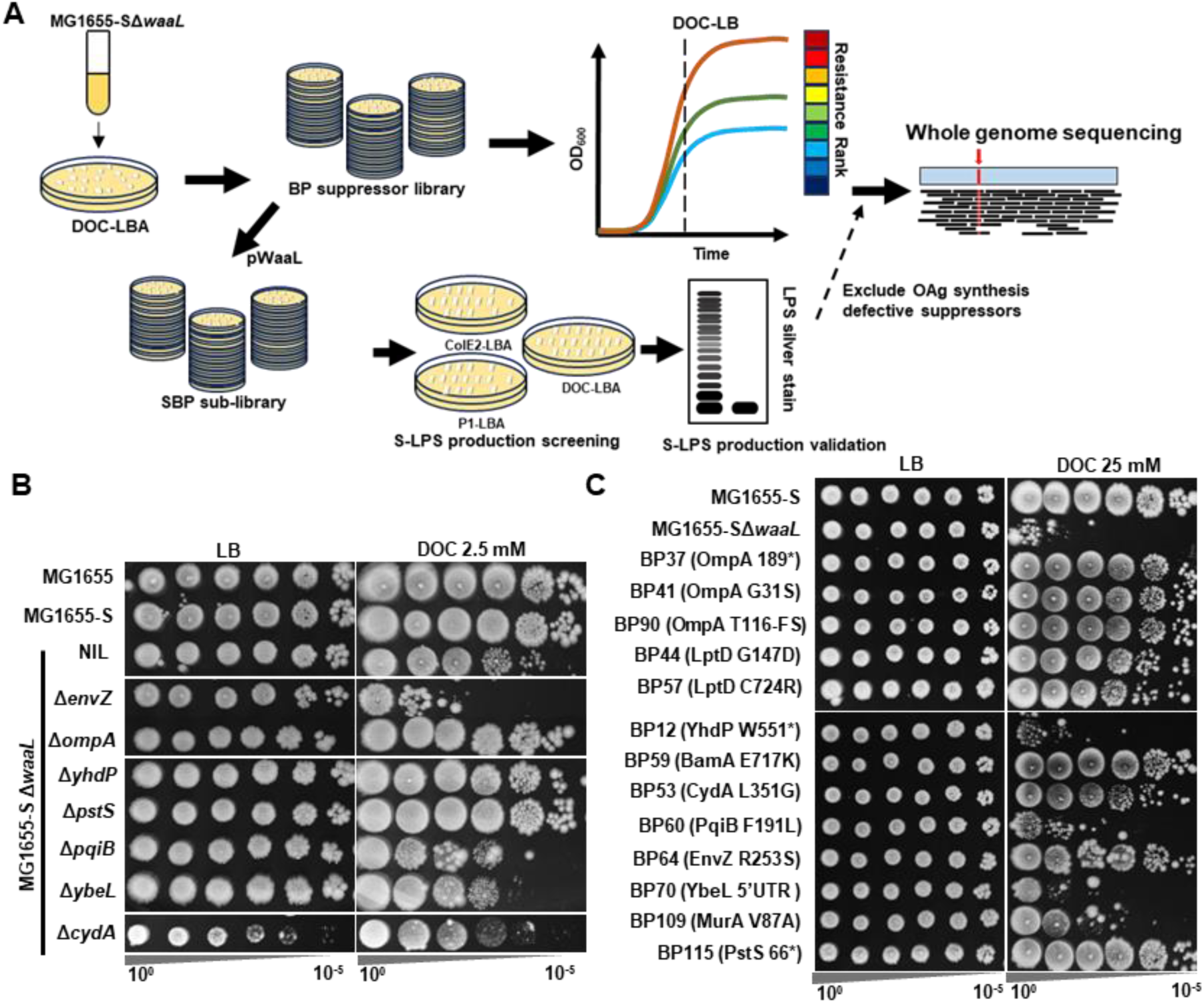
Isolation, selection and characterisation of MG1655-SΔ*waaL* BS-resistant suppressor mutants. **(A)** Schematic representation of suppressor mutant isolation and selection for whole genome sequencing. See Methods and Materials for detailed description. **(B)** BS-sensitivity assay for the validation of restored BS resistance of suppressor mutations identified targeting non-essential genes through single gene deletion mutants. **(C)** Selection of strong suppressor by BS sensitivity assay with elevated concentration of BS (25 mM DOC). FS, frameshift mutation.

Interestingly, two SBP mutants (SBP66 and SBP73) produced only the OAg-deficient LPS (rough-LPS, R-LPS) with a single OAg RU (termed semi-rough-LPS, SR-LPS) (S2A Fig), characteristic of a MG1655-SΔ*wzyB* mutant [3], suggesting that their corresponding BP mutants might have a *waaL-wzyB-null* genotype. However, we have previously shown that disruption of both *waaL* and *wzyB* is lethal in MG1655-S as a result of the complete stall of UndPP-OAg in the periplasm [21]. We therefore performed whole genome sequencing (WGS) for these two mutant isolates and identified mutations in *wecA* and *wbbL* (Table 1), whose protein products are the initial and second glycosyltransferases (IT and 2^nd^ GT) for the assembly of OAg RU, thus responsible for the engaging and the committed steps of OAg RU assembly, respectively [21]. We reasoned that these mutations (predicted amino acid substitutions) might result in reduced enzymatic activities of WecA and WbbL, leading to a reduced cellular UndPP-OAg, inadequate for WzyB polymerisation and outcompeted by WaaL in the periplasm [22] to give a MG1655-SΔ*wzyB*-like LPS profile. These results confirmed our model that the prolonged accumulation of UndPP-OAg intermediates sensitises *E. coli* to BS [3], and this sensitisation is strongly suppressed by mutations in the OAg biosynthesis pathways.

**Table 1.** Mutations identified in MG1655-SΔ*waaL* suppressors grown in DOC-LBA.

| BP Mutants | Nucleotide Mutation and locus | Peptide mutation | LPS in SBP |
| --- | --- | --- | --- |
| BP12 | C-T substitution at 3394727 | YhdP W551* | S-LPS |
| BP37 | G-C substitution at 1019487 | OmpA Y189* | S-LPS |
| BP41 | C-T substitution at 1019963 | OmpA G31S | S-LPS |
| BP44 | C-T substitution at 56670 | LptD G147D | S-LPS |
| BP53 | T-C substitution at 772509 | CydA L351G | S-LPS |
| BP57 | A-G substitution 54940 | LptD C724R | S-LPS |
| BP59 | G-A substitution at 200552 | BamA E717K | S-LPS |
| BP60 | C-A substitution at 1013831 | PqiB F191L | S-LPS |
| BP64 | G-T substitution at 3535112 | EnvZ R253S | S-LPS |
| BP70 | G-A substitution at 6774853 | 5'UTR of *ybeL* | S-LPS |
| BP90 | Single A deletion at 1019706 | OmpA frameshift at T116 | S-LPS |
| BP109 | A-G substitution at 3336235 | MurA V87A | S-LPS |
| BP115 | Single C deletion at 3911349 | PstS truncated to 66 AA | S-LPS |
| BP66 | C-G substitution at 3974971 | WecA P272R | SR-LPS |
| BP73 | T-C substitution at 2104434 | WbbL Y3C | SR-LPS |

### Disruptions in OM biogenesis machineries restored BS resistance in MG1655-SΔwaaL

In the top 31 ranked BS resistant BP mutants (S1 Table), we excluded 11 with mutations potentially affecting OAg synthesis. The remaining 20 BP mutants were confirmed to have restored resistance to BS in comparison to MG1655-SΔ*waaL* (S2C Fig). Interestingly, 8 BP mutants (BP12, BP41, BP44, BP57, BP59, BP60, BP70 and BP90) lacked the elevated vancomycin resistance of MG1655-SΔ*waaL* [3] (S2C Fig). These 8 BP mutants along with another 5 BP mutants (BP37, BP53, BP64, BP109 and BP115), which maintained the elevated vancomycin resistance (S2C Fig), were further confirmed to have no detectable changes in S-LPS profile when complemented by WaaL in the SBP sub-library (S2D Fig) and were subjected to WGS analysis (Fig 2A).

WGS identified 8 mutations affecting different OM components and 5 mutations in other genes. We found mutations in OM-related genes encoding one of the most abundant outer membrane proteins (OMP), OmpA (BP37, BP41 and BP90), an essential component of the OMP biogenesis machinery, BamA (BP59), an essential component for LPS translocation, LptD (BP44 and BP57), and two proteins (YhdP and PqiB) involved in the lipid transport to the OM (BP12 and BP60) (S2C Fig & Table 1). We also identified 5 additional mutations in genes encoding the following: the sensor histidine kinase EnvZ (BP64), which modulates OM porins OmpC and OmpF; cytochrome bd-I ubiquinol oxidase subunit 1 CydA (BP53), which acts as a terminal oxidase producing proton motive force; UDP-N-acetylglucosamine enolpyruvoyl transferase MurA (BP109), which catalyses the first committed step in the assembly of PG; the periplasmic phosphate binding protein PstS (BP115); the 5’UTR of the protein of unknown function, YbeL (BP70) (S2C Fig & Table 1).

To validate these suppressor mutations, we generated single gene deletion mutants of the non-essential genes *envZ*, *ompA*, *yhdP*, *pstS*, *pqiB*, *ybeL* and *cydA* in the MG1655-SΔ*waaL* strain background and examined their resistance to BS. We confirmed that deletions of *ompA*, *yhdP* and *pstS* restored BS resistance in MG1655-SΔ*waaL* (Fig 2B), suggesting that mutational alterations identified in these genes likely resulted in a loss of function (Fig 2B). In contrast, deletions in *pqiB, ybeL* and envZ did not rescue BS resistance of MG1655-SΔ*waaL* (Fig 2B), with deletion of *envZ* instead further sensitised MG1655-SΔ*waaL* to BS (Fig 2B), suggesting that mutational alterations identified in these genes might lead to altered protein activity. The effect of *cydA* deletion on BS resistance of MG1655-SΔ*waaL* could not be determined as the growth of MG1655-SΔ*waaL*Δ*cydA* in plain LBA was compromised (Fig 2B). These results confirmed our genotypic analysis and suggested complex, multiple pathways for rescuing BS resistance of MG1655-SΔ*waaL*.

### Screening for strong suppressors

To further select strong suppressor mutations that confer BS resistance of MG1655-SΔ*waaL*, we challenged the selected 13 BP suppressor mutants with an elevated concentration of BS (25 mM DOC) and found that among all tested suppressors, the mutations identified in *ompA, lptD, bamA, cydA, envZ* and *pstS* conferred the highest BS resistance to MG1655-SΔ*waaL* (Fig 2C). An EnvZ^P41L^ mutant has previously been shown to influence BS resistance [23] through altering expression levels of OmpC and OmpF [24]. OmpF was proposed to be a porin for BS OM entry [25] and OmpC was shown to be required for BS resistance [26]. Indeed, deletion of *ompC* slightly further sensitised MG1655-SΔ*waaL* to BS, while deletion of *ompF* slightly improved its BS resistance (Fig 3A). We next examined OmpC and OmpF levels in all of the strong suppressor mutants, which revealed that the mutants with BamA^E717K^ or EnvZ^R253S^ have reduced level of OmpF/C (Fig 3B), while other suppressor mutants had no detectable changes in the levels of OmpF/OmpC.

**Fig 3.**
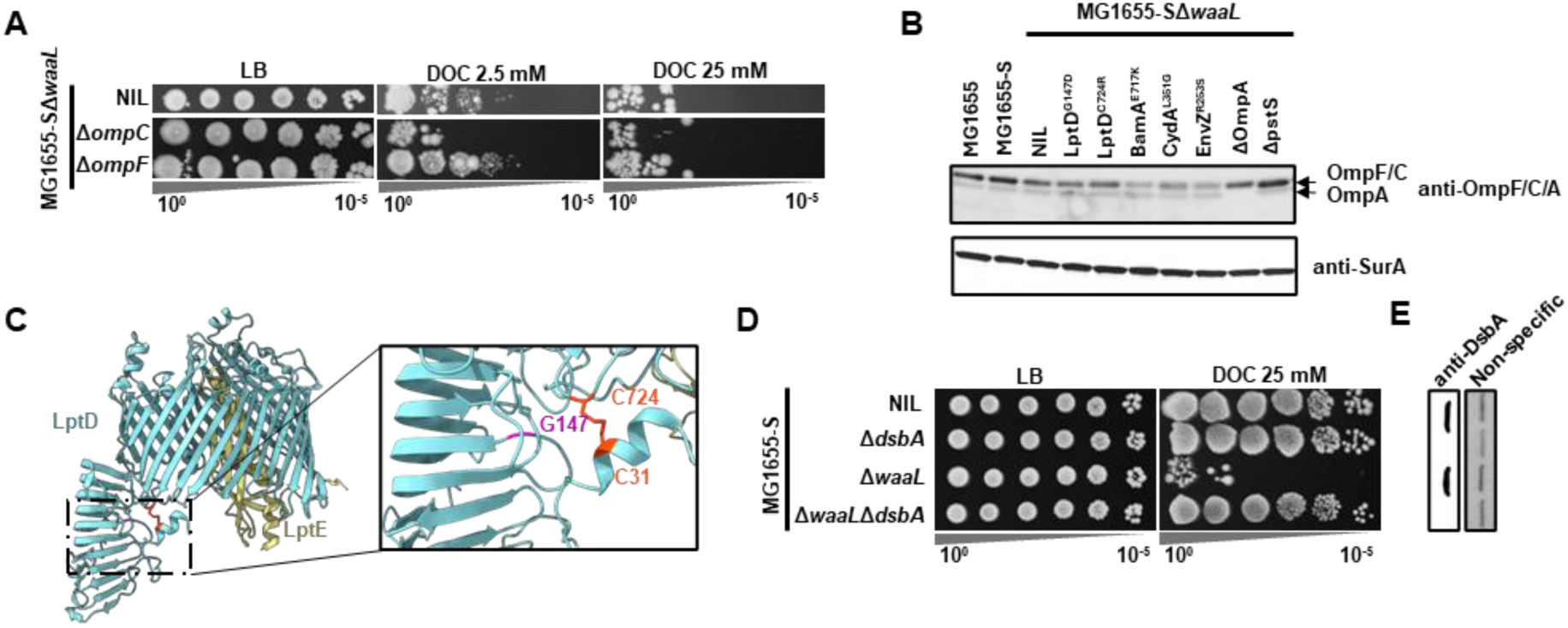
Investigation of strong MG1655-SΔ*waaL* BS suppressor mutations. (**A)** BS sensitivity assay of single gene deletion mutants in MG1655-SΔ*waaL* with 2.5 mM or 25 mM DOC. (**B**) Western immunoblotting of OmpF, OmpC and OmpA from whole cell lysate prepared from indicated bacterial strains with anti-OmpF/C/A antibodies. Anti-SurA antibody was used to detect SurA expression as a loading control. **(C)** Identified LptD suppressor mutation mapping on the crystal structure of LptD from *Shigella flexneri* (PDB 4Q35). Mutated residues G147 and C724 are shown in magenta and orange, respectively, with the C31-C724 disulfide bridge shown. **(D)** BS sensitivity assay of single gene deletion mutants in MG1655-SΔ*waaL* with 2.5 mM or 25 mM DOC. **(E)** Western immunoblotting of DsbA from whole cell lysate prepared from indicated bacterial strains with anti-DsbA antibodies, and a non-specific protein band detected with anti-DsbA was used to indicate sample loading.

LptD is an essential component of the LPS transport machinery across the OM. It features a two-domain structure: a barrel that is embedded in the OM and a periplasmic domain that structurally resembles the LptA subunit. This folding enables LptD to effectively connect with periplasmic LptA to form the transenvelope bridge for LPS transport [27]. We mapped two independent mutations located at the interdomain interface (Fig 3C) both with reduced resistance to vancomycin (S2C Fig) yet without detectable differences in S-LPS production (S2C Fig) and ColE2 resistance (S1 Table) when complemented with WaaL. Interestingly, one of the suppressor mutants, LptD^C724R^, abolishes one of the two known disulfide bridge (C31-C724) in LptD [28]. Although the physiological importance of the disulfide bridge between the two domains of LptD is unclear, it was shown to be catalysed by the periplasmic oxidative foldase DsbA [28]. Consistent with the known role of DsbA in LptD disulfide bridge formation, disruption of *dsbA* in MG1655-SΔ*waaL* fully restored its BS resistance (Fig 3D-E), however we were unable to determine whether the restoration of BS resistance observed in MG1655-SΔ*waaL*Δ*dsbA* was solely due to the compromised disulfide bond formation in LptD.

### Characterisation of pstS and bamA mutations that restore BS resistance in MG1655-SΔwaaL

Disruption of *pstS* was shown previously to up-regulate the expression of UgpB bearing moonlighting function as a periplasmic chaperone that aids protein folding to confer BS resistance [29]. However, in MG1655-SΔ*waaL*: i) ectopic expression of UgpB did not restore BS resistance, ii) disruption of *ugpB* did not further sensitise the strain to BS, and iii) disruption of *ugpB* in the suppressor mutant MG1655-SΔ*waaL*Δ*pstS* did not re-sensitise it to BS either (S3C Fig), suggesting that the rescue of BS resistance of MG1655-SΔ*waaL* by the disruption of *pstS* is not through UgpB chaperone function.

The BS DOC was reported previously as a potent protein-unfolding agent posing disulfide stress [4], and a periplasmic chaperone FkpA was strongly upregulated in a *pstS* deletion mutant [29]. We therefore investigated the role of several selected periplasmic chaperones (DegP, Skp, SurA, FkpA and Spy) in BS resistance of MG1655-SΔ*waaL*. Interestingly, none of these chaperones was required in maintaining the BS resistance of MG1655-SΔ*waaL* as disruption of neither of them further sensitises MG1655-SΔ*waaL* to BS (Fig 4A). However, counterintuitively, disruptions of *surA*, *fkpA* and *spy* instead rescued BS resistance (DOC 2.5 mM) in MG1655-SΔ*waaL,* with disruption of *surA* in MG1655-SΔ*waaL* conferring the highest BS resistance (DOC 25 mM), to the same level to that of MG1655-SΔ*waaL*Δ*ompA* (Fig 4A). Additionally, MG1655-SΔ*waaL*Δ*surA* was found to have reduced OmpA protein levels, with a high protein level of DegP detected (Fig 4B), indicating a severe OM protein folding stress.

**Fig 4.**
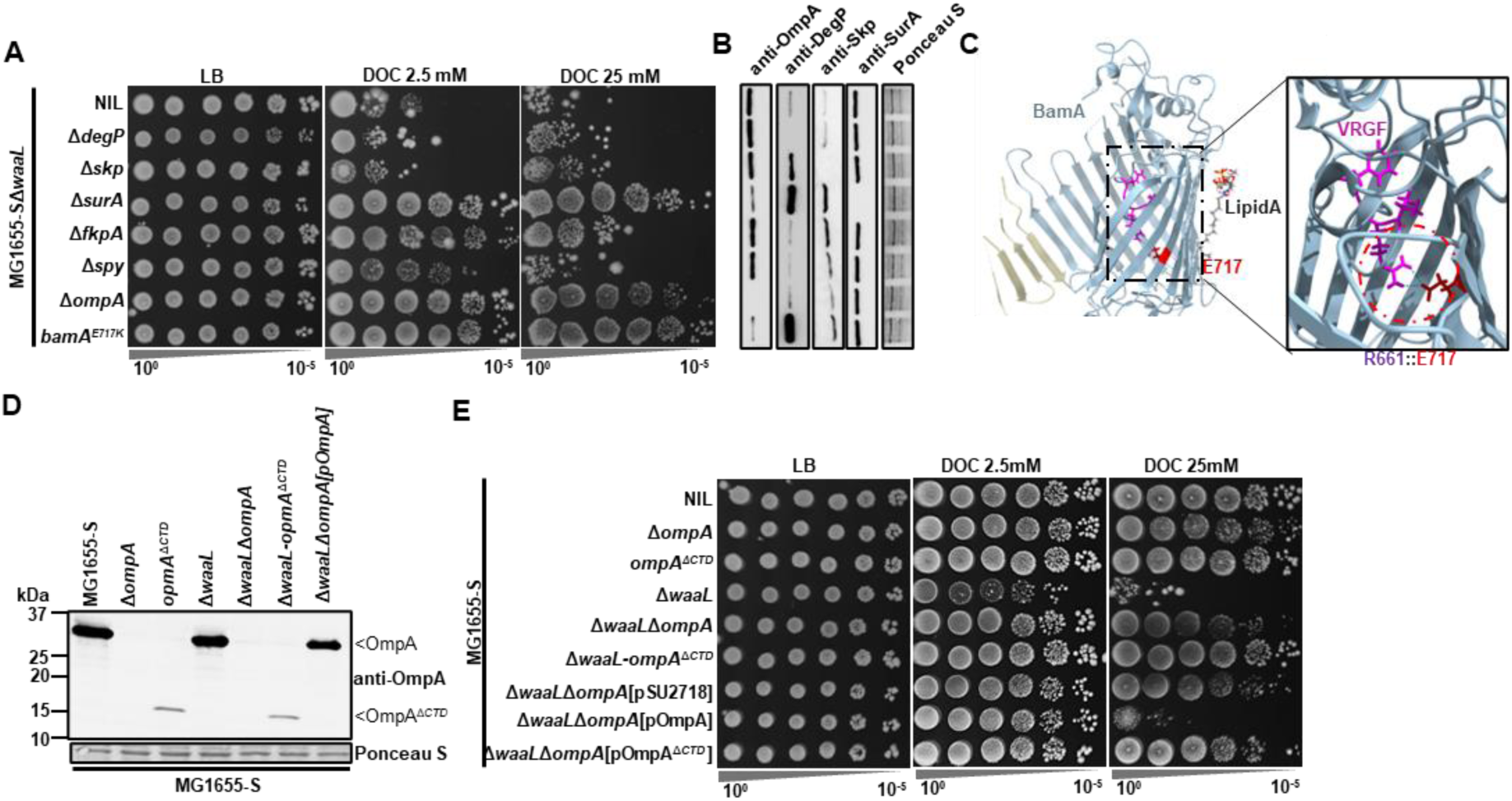
Disruption of OmpA CTD confers BS resistance to *MG1655-S*Δ*waaL*. (**A)** BS sensitivity assay of single gene deletion mutants in MG1655-SΔ*waaL* with 2.5 mM or 25 mM DOC. (**B**) Western immunoblotting for detection of OmpA, DegP, Skp, SurA from whole cell lysate prepared from indicated bacterial strains with anti-OmpA, anti-DegP, anti-Skp and anti-SurA antibodies. Transferred proteins from cell lysate were stained with Ponceau S to indicate sample loading. The molecular weight of protein markers is indicated on the left to validate the size of the truncated OmpA protein OmpA^ΔCTD^. (**C**) BamA suppressor mutation mapping on its barrel structure (PDB 7TTC). The salt bridge between E717 and R661 in the conserved VRGF motif is shown. (**D**) Western immunoblotting for detection of OmpA from whole cell lysate prepared from indicated bacterial strains with anti-OmpA antibodies. Transferred proteins from cell lysate were stained with Ponceau S to indicate sample loading. **(E)** BS sensitivity assay of single gene deletion mutants in MG1655-SΔ*waaL* with 2.5 mM or 25 mM DOC.

SurA is important in OMP folding as it delivers its OMP clients to BamA for folding into the OM [30]. In the suppressor mutant BP59, a mutation E717K was mapped on the OMP foldase BamA of the BAM foldase structure (Table 1). BamA E717 forms salt bridges with R661 in BamA which is important in stabilising a conserved VRGF motif (Fig 4C) [31]. Disruptions of the VRGF motif was shown previously to compromise BamA folding activity, leading to reduced levels of OMPs, including OmpF, OmpC and OmpA [31]. Indeed, the BamA^E717K^ suppressor mutant in the MG1655-SΔ*waaL* background also showed reduced levels of OmpF, OmpC (Fig 3B) and OmpA (Fig 4B). Intriguingly, in the 13 selected suppressors conferring top-ranked BS resistance to MG1655-SΔ*waaL,* 3 independent mutations were mapped to affect OmpA and these conferred the highest BS resistance (Table 1). Additionally, the BamA^E717K^ mutation and *surA* deletion were also found with reduced OmpA levels conferring high BS resistance to MG1655-SΔ*waaL,* although we were unable to determine whether the rescue effect was due to reduced OmpA level alone. Together, these results suggest that the biogenesis of OmpA poses stress in MG1655-SΔ*waaL* growing in BS, and we therefore focused our further studies on *ompA* mutations.

### Disruptions of OmpA CTD restores BS resistance in MG1655-SΔwaaL

OmpA is predicted to be a two-domain protein with an N-terminal domain (NTD, aa 22-192) as an 8-strand beta-barrel embedded in the OM, and a C-terminal domain (CTD, aa 193-346) folded independently in the periplasm. Interestingly, the mutation found in the BP37 would truncate the CTD of OmpA but allow its barrel to be produced. We therefore generated chromosomal deletions of OmpA CTD (aa Δ190-346, designate as OmpA^ΔCTD^) in both MG1655-S and MG1655-SΔ*waaL* and confirmed the truncation of OmpA, which was detected at reduced levels (Fig 4D). While both *ompA* and *omp*^Δ*CTD*^ had no detectable effect on cell survival of MG1655-S in BS containing media (Fig 4E), these mutations increased BS resistance in MG1655-SΔ*waaL*. Intriguingly, *omp*^Δ*CTD*^ conferred greater BS resistance in comparison to Δ*ompA* in MG1655-SΔ*waaL* (Fig 4E). This was further confirmed via the OmpA complementation experiment showing that while the complementation of OmpA in the MG1655-SΔ*waaL*Δ*ompA* mutant re-sensitised the strain to BS, complementation of OmpA^ΔCTD^ did not (Fig 4E). Together, these results strongly suggest that the production of OmpA barrel is beneficial to BS resistance while the production of OmpA with its CTD is detrimental to the survival of MG1655-SΔ*waaL* in the presence of BS.

### OM-PG anchoring is detrimental to BS resistance in MG1655-SΔwaaL

OmpA CTD has previously been shown to be involved in the regulation of Rcs stress response through interacting with RcsF [32]. However, we found that the disruption of *rcsF* had no effect in rescuing the BS resistance in MG1655-SΔ*waaL* (S4 Fig). The periplasmic CTD of OmpA adopts a globular structure that is highly similar to known PG-binding domains of RmpM (*Neisseria meningitidis*) and MotB (*Helicobacter pylori*) [33] and has been proposed to interact with PG [34]. Therefore, it is possible that the OM tethering through interaction between OmpA and PG sensitises MG1655-SΔ*waaL* to BS.

To test this hypothesis, we investigated another abundant protein that tethers the OM and PG is Braun’s lipoprotein (Lpp), which is covalently linked to PG by three redundant LD-transpeptidases, LdtA, LdtB (catalyses majority of the Lpp-PG attachment) and LdtC, between the Lpp^K58^ and PG^mDAP^ residues [35]. We therefore next deleted *lpp* in MG1655-SΔ*waaL*. Surprisingly, disruption of *lpp* fully restored BS resistance of MG1655-SΔ*waaL* (Fig 5A-B), and the restoration of BS resistance was due to the PG-Lpp linkage, since disruptions of *ltdA*, *ltdB*, or *ltdC* all marginally restored BS resistance (Fig 5A) and the point mutant of Lpp^ΔK58^ fully restored BS resistance (Fig 5A). The linkage between Lpp and PG is lethal when lipoprotein transport is impaired, leading to the PG-attachment of IM-accumulated Lpp [36]. To investigate whether BS affected lipoprotein transport in MG1655-SΔ*waaL*, we fractionated membranes of MG1655, MG1655-S and MG1655-SΔ*waaL* grown in the presence and absence of BS (S5A Fig), and confirmed the absence of Lpp in the IM fractions and the successful separation of IM and OM of all strains growing in the presence of BS (S5B Fig). These results suggest that the lipoprotein transport was not affected in MG1655-SΔ*waaL* when growing in the presence of BS. Additionally, we also confirmed that UndPP-OAg still accumulated when growing in the presence or absence of BS for all strong suppressor mutants reported in this work, including MG1655-SΔ*waaL*Δ*lpp* (S6 Fig), indicating that the restoration of BS resistance in these strong suppressor mutants was not due to the loss of OAg intermediate accumulation. Taken together, these results suggest that abundant OM-PG tethering renders MG1655-SΔ*waaL* unfit for growth in the presence of BS.

**Fig 5.**
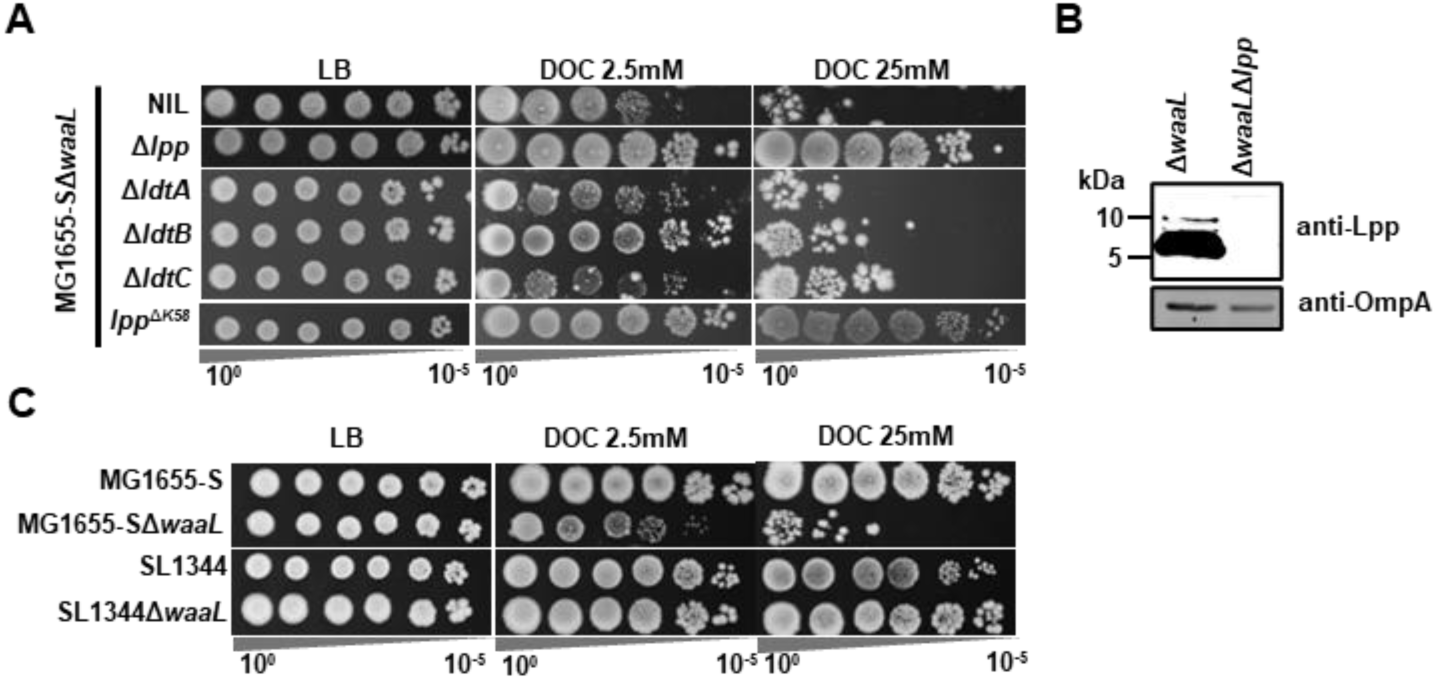
Disruption of Lpp and PG linkage confer BS resistance to *MG1655-S*Δ*waaL*. **(A)** BS sensitivity assay of single gene deletion mutants and protein mutants in MG1655-SΔ*waaL* with 2.5 mM or 25 mM DOC. (**B**) Western immunoblotting of Lpp from whole cell lysate prepared from indicated bacterial strains with anti-Lpp antibodies. OmpA was detected with anti-OmpA antibodies to serve as a loading control. Molecular weight is indicated. **(C)** BS sensitivity assay indicated strains with 2.5 mM or 25 mM DOC.

BS was shown previously to induce PG remodelling with reduced level of Lpp-PG linkage in *Salmonella enterica* SL1344 [37]. However, MG1655-SΔ*waaL* growing in the presence and absence of BS (S3 Table & S7 Fig) resulted in similar level of Lpp-PG crosslinking. Interestingly, unlike MG1655-SΔ*waaL*, *S. enterica* SL1344Δ*waaL* was not hyper-sensitive to BS (Fig 5C). These results further indicate that loss of crosslinking between OM and PG confers BS resistance.

## Discussion

### Accumulation of UndPP-OAg is a major contributor to the sensitivity to BS

In this study, we compared the BS sensitivity across a set of isogenic mutants in OAg-producing *E. coli* K-12 strain, and revealed that all core-oligosaccharide truncation mutants accumulated UndPP-OAg and had elevated BS sensitivity in comparison to their corresponding OAg-defective strains. This work strongly supports our model that the presence and accumulation of polymerised UndPP-OAg in the periplasm poses stresses when growing in the presence of BS [3]. Mutations affecting LPS inner-core oligosaccharide synthesis, termed deep-rough LPS mutants, especially the heptoseless mutants, were reported to be more sensitive to hydrophobic but not to hydrophilic antibiotics [38]. One might argue that the increased sensitivity to BS was due to compromised OM permeability as the results of loss phosphate substitution that facilitate LPS-Ca^2+^ bridges. However, we showed that the chelation rather than addition of Ca^2+^ in the culture media improved the resistance of MG1655-SΔ*waaL* to BS, suggesting that the addition of Ca^2+^ adversely affects membrane integrity in this situation and hence lateral LPS-LPS interaction through Ca^2+^ bridging plays a limited role in conferring BS resistance. The deep-rough LPS mutants have a reduced level of OMPs and disturbed OM asymmetry due to patches of phospholipids on the outer leaflet [17, 39]. It is possible that such phospholipid bilayer patches in the deep-rough LPS allow the passage of BS. Nevertheless, our results suggest that the accumulation of UndPP-OAg is a major contributor to the sensitivity to BS.

### Compensatory role of OM phospholipid architecture in conferring BS resistance

Since the sensitisation to BS in strains with accumulated periplasmic UndPP-OAg did not appear to affect efflux pump activity, we further investigated the mechanisms by which BS inhibited the growth of strains that accumulate OAg intermediates in the periplasm. Characterisation of a set of suppressor mutants surviving a lethal dose of BS identified, as expected, suppressors with abolished OAg biogenesis. Unexpectedly, we also identified suppressors with compromised OMP biogenesis amongst isolates with the highest level of BS resistance. One of these suppressor mutations affects the anterograde phospholipid transport component YhdP, which is predicted to have a trans-envelope structure spanning between IM and OM with a hydrophobic groove for phospholipid binding [40] (Fig 6). Similarly, a substitution mutation in the protein PqiB with a tube-like trans-envelope structure was also found as a suppressor. PqiB belongs to the mammalian cell entry (MCE) superfamily, whose protein members were also implicated in lipid transport [41]. Both proteins were reported previously to affect the OM [42, 43]. We speculate that the mutations facilitate the formation of phospholipid patches on the outer leaflet of the OM, which appear to be beneficial in MG1655-SΔ*waaL*, explaining the reduced vancomycin resistance of the suppressors compared to their parent strain. That these suppressor mutants could not withstand elevated concentrations of BS further supports a compensatory role through changing the OM phospholipid architecture.

**Fig 6.**
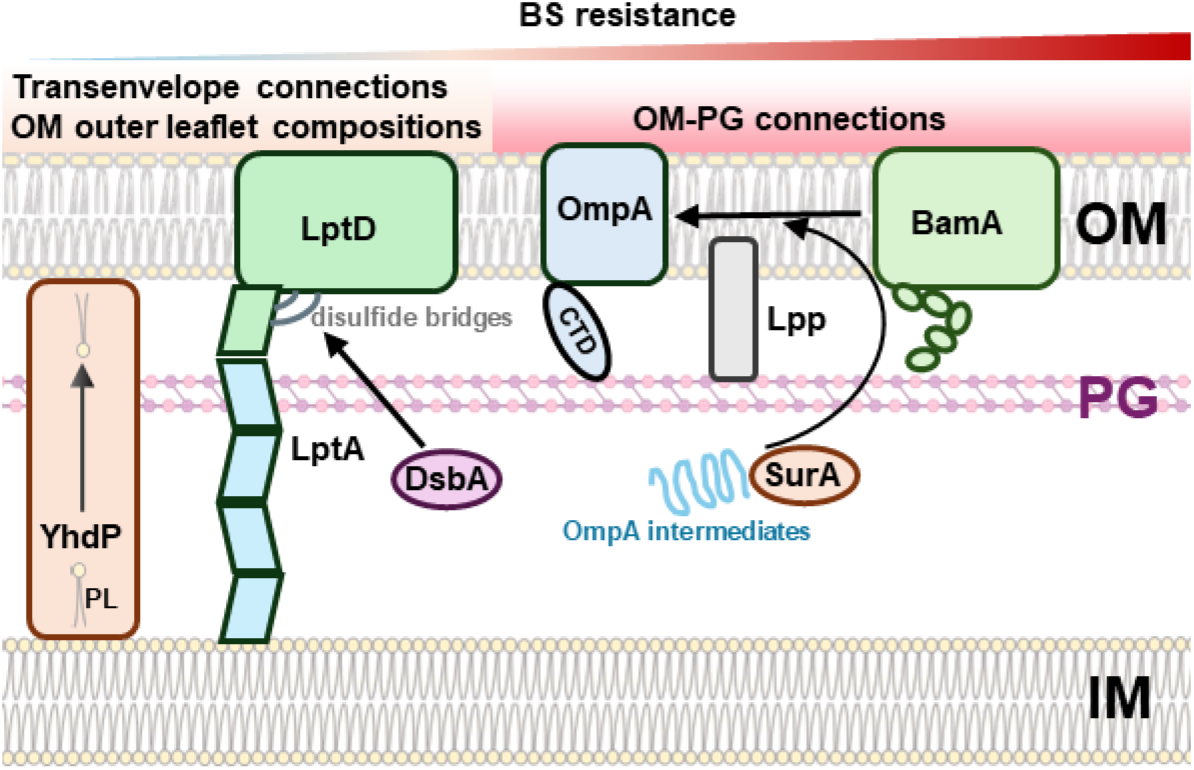
Summary of major cell envelope suppressor mutants with restored BS resistance in MG1655-SΔ*waaL*. UndPP-OAg poses cell envelope stresses in the presence of BS, presumably through affecting PG biosynthesis. Disruptions in YhdP which is responsible for phospholipid (PL) transport confer weak BS resistance. Disruptions in LptD interdomain interactions, OM-PG connecting molecules OmpA and Lpp, OMP biogenesis components BamA and SurA, as well as cell envelope oxidase/foldase DsbA confer strong BS resistance.

### Weakened LptD interdomain interactions confer high BS resistance

At the elevated concentration of BS, we acquired two independent mutants with suppressor mutations mapped in LptD, the two-domain OM protein responsible for the translocation of LPS molecules (Fig 6). The interdomain of LptD forms two pair of disulfide bridges by four non-consecutive cysteine residues [28], which are dependent on the periplasmic protein oxidase/foldase DsbA [28]. Consistent with these observations, we showed that disruption of *dsbA* in MG1655-SΔ*waaL* also fully rescued its BS resistance, similar to the two *lptD* suppressor mutations. This seems also to be supported by the suppressor mutation found in *cydA*. CydA is a subunit of cytochrome *bd*, the quinol oxidase that participates in the bioenergetics of bacterial cell, providing the source of oxidising power which fuels the DsbA-DsbB-ubiquinone complex for the catalysation of disulfide bridges in folding periplasmic proteins [44]. It is likely that the suppressor mutations affecting CydA also indirectly affect LptD intramolecular interdomain disulfide bond formation. The other suppressor mutation (LptD^G1457D^) was also found in the interdomain interface. These two LptD suppressor mutants all exhibited reduced level of vancomycin resistance in comparison to MG1655-SΔ*waaL*, again suggesting that the rescue was not through excluding BS via improved OM barrier function. Additionally, when complemented with pWaaL, these two mutants were able to produce S-LPS in a level indistinguishable from pWaaL complemented MG1655-SΔ*waaL* via LPS silver staining and colicin E2 sensitivity assay, indicating that the production and translocation of S-LPS might not be affected. It thus remains unclear what role of LptD was affected when interdomain interaction is altered through these two suppressor mutations. One possibility is that the interactions between LptD barrel and its periplasmic domain connecting to LptA limits the tethering angle of the trans-envelope LptA bridge, thereby conferring regional membrane stiffness. The disruption of these connections would likely increase the regional OM-PG-IM flexibility, resulting in increased BS resistance. However, this is beyond the scope of this study and warrants future research with defined mutations to investigate the role of LptD interdomain interactions. Nevertheless, our data tempt us to suggest that altered/weakened LptD interdomain interaction could confer MG1655-SΔ*waaL* BS resistance.

### Compromised OMP biogenesis machinery confer high BS resistance

In attempting to search for the potential role of other well-known periplasmic chaperones in conferring BS resistance in MG1655-SΔ*waaL,* we unexpectedly found that the disruption of *surA* conferred full resistance of BS in MG1655-SΔ*waaL.* SurA is an important chaperone that directs client OMPs to BamA for their assembly on the OM [30] (Fig 6). Both the loss of SurA and destabilisation of conserved motif (VRGF) of BamA fully restored BS resistance in MG1655-SΔ*waaL* presumably through a reduced production of OMP including OmpA, which was found with three independently mapped suppressor mutations. OmpA is a key cell envelope protein that provides tensile strength of the cell envelope, with it barrel domain critical for maintaining the barrier function of OM [45]. Indeed, OmpA was previously linked in both OAg-producing *E. coli* O157 [46] and in non-OAg *E. coli* K-12 [45] to be required for BS resistance. On the contrary, we found that loss of OmpA restored BS resistance of MG1655-SΔ*waaL* to the level of MG1655-S. Surprisingly, it was the C-terminal periplasmic domain of OmpA that sensitised MG1655-SΔ*waaL* to BS, while the possession of the N-terminal porin domain was beneficial in conferring greater BS resistance in comparison to the complete loss of OmpA protein.

### Weakened the interactions between OM and PG confer high BS resistance

The C-terminal domain of OmpA interacts with PG, contributing to the tethering of the OM to the PG [34] (Fig 6). Our data suggests a model that normal-level OM tethering to PG layer reduces the fitness of cells exposed to BS and with OAg intermediates accumulating in the periplasm. Consistent with this model, the disruption of *lpp*, encoding the most abundant OM lipoprotein that covalently anchors to PG (Fig 6), likewise fully restored BS resistance of MG1655-SΔ*waaL*. Moreover, the individual disruptions of functionally redundant transpeptidase genes (*ldtABC*) increased the BS resistance of MG1655-SΔ*waaL* and mutating the residue of Lpp responsible for the PG-attachment completely restored BS resistance of MG1655-SΔ*waaL.* This is consistent with a previous report in *S. enterica* where disruptions of transpeptidases catalysing the covalent attachment of Lpp to PG resulted in hyper-resistance to BS [37]. However, unlike *E. coli* K-12 MG1655-SΔ*waaL*, *S. enterica* SL1344Δ*waaL* was found to be fully resistant to BS. In line with our model, this could be explained by a reduced level of Lpp-PG linkages that were reported for *S. enterica* cells growing in the presence of BS [37], whereas MG1655-SΔ*waaL* showed similar levels of Lpp-PG linkages in the presence and absence of BS. The drastic differences may be accounted for by the unique human niche of *S. enterica* compared to other BS-resistant enteric bacteria, in which the species exhibits a striking ability to withstand exceptionally high concentrations of BS (>60%) to allow their survival in the gallbladder, the storage site for bile [47]. In hindsight of this work, *S. enterica* might be able to remodel its Lpp-PG linkages in response to BS, an adaptive regulatory mechanism that appears to be absent in *E. coli* K-12. Hence, *Salmonella spp*.’s ability to alter the degree of Lpp-PG crosslinking in response to BS exposure might contribute to its adaptation to host niches with high bile concentrations, setting them apart from other enteric bacteria.

Another trans-envelope molecular complex that known to connect OM to PG is the Tol-Pal system. Likewise, the natural occurring intragenic tandem repeats variation found in the coding regions of periplasmic component TolA with coiled structure was reported to play a role in modulating BS resistance, with higher number of tandem repeats giving a longer TolA in molecular length conferring higher BS resistance [48]. Extension in the molecular length of TolA molecules may provide greater flexibility to OM, with increased membrane fluidity. This seems to agree with the data generated from OmpA suppressor mutants. A recent study reported the connection between OM to PG with OmpA was found to maintain tensile strength and OM stiffness [45]. This predicts that the loss of OmpA would increase the fluidity of OM. Collectively these lines of evidence suggest that loss/weakening the interactions between OM and PG may decrease membrane stiffness when OAg is produced, conferring cells with benefits in adapting to high concentrations of BS.

In conclusion, this study provides compelling evidence that a reduction in the strength of OM-PG interactions in the OAg-producing bacteria significantly enhances resistance to BS. This strategy likely enables enteric bacteria to adapt to host niches where BS are present. Our findings highlight the critical role of cell envelope plasticity in enabling bacterial adaptation to challenging host environments.

## Materials and Methods

### Bacterial strains and plasmids

The bacterial strains and plasmids used in this work are listed in S2 Table. Single colonies of bacterial strains grown overnight on Lysogeny Broth (LB)-Lennox [49] agar (1.5% w/v) plates were picked and grown overnight in LB at 37 °C for subsequent experiments. Where appropriate, media were supplemented with ampicillin (Amp, 100 µg/ml), kanamycin (Kan, 50 µg/ml), chloramphenicol (Chl, 25 µg/ml), vancomycin (Van, 200 µg/ml), sodium deoxycholate (DOC, 2.5 mM or 25 mM), calcium chloride (Ca^2+^, 5 mM), Ethyleneglycol-bis(β-aminoethyl)-N,N,Nʹ,Nʹ-tetraacetic Acid (EGTA, 1 mM), purified Colicin E2 DNA endonuclease (ColE2, 100 μg/ml) [3], anhydrotetracycline (AhTet 50 ng/ml), or arabinose (Ara, 10 mM).

### Bacterial phage and phage lysate preparation

Bacteriophage preparation was done as described previously [3]. Briefly, bacteriophage P1kc (ATCC 11303-B23) lysate was prepared by combining 100 μl of mid-exponential culture of MG1655 with phage stocks at a multiplicity of infection (MOI) of approximately 0.5 in 3 ml of LB soft agar (0.75% w/v agar) supplemented with MC salts (100 mM MgSO_4_ and 5 mM CaCl_2_). This mixture was then poured onto LB agar plates and incubated at 37 °C for 18 h. The clear upper layer of soft agar containing the phage lysate was carefully scraped off, mixed with 2 ml of LB-Miller media (LB supplemented with MC salts), vortexed, and centrifuged. The resulting clear supernatant, which contained the bacteriophage, was collected and sterilised by adding 10 μl of chloroform. Phage titres were determined by spotting 5 μl of tenfold serially diluted phage stock onto top LB soft agar plates inoculated with 100 μl of MG1655 mid-exponential culture. Plaque-forming units (pfu) were counted after incubating the infected plates at 37 °C for 18 h.

### Plasmid construction

For generation of expression constructs, the coding sequences of targeted proteins were PCR amplified from genomic DNA prepared via Qiagen QIAamp DNA Blood Mini Kit per manufacturer’s protocol and cloned into indicated plasmids using restriction enzyme cloning (NEB).

### Bacterial mutagenesis via allelic exchange

Mutagenesis was conducted following previously established protocols [50] with laboratory-optimised modifications [51]. Briefly, bacterial strains containing the plasmid pKD46 were cultured overnight in 10 ml of Luria-Bertani (LB) broth at 30 °C and subsequently sub-cultured at a ratio of 1:100 into 10 ml of LB in a 50 ml tube. Induction of lambda phage-derived protein expression was achieved by adding 50 mM L-arabinose when the optical density at 600 nm (OD600) reached 0.3, followed by a 1-h incubation. The bacterial cells were harvested through centrifugation, washed twice with 10 ml of ice-cold water, and resuspended in 100 μl of 10% (v/v) ice-cold glycerol for electroporation. The *cat* or *neo* gene was amplified by PCR from pKD3 or pKD4, respectively, using primers that included 40-50 bp of homologous sequences flanking the target gene (S2 Table). The purified PCR amplicon (1.5 μg) was introduced into electrocompetent cells via electroporation, and the cells were immediately allowed to recover in 3 ml of LB in a 50 ml Falcon tube for 2 h at 37 °C before plating 100 μl onto LB agar plates supplemented with chloramphenicol (Chl) or kanamycin (Kan). The plates were then incubated at 37 °C for 16 h to facilitate the selection of mutants. Successful mutants were subsequently screened and confirmed by PCR.

All mutagenesis including both knocking out and knocking in were done with lambda red mutagenesis and the primers were listed in S2 Table with clear annotations. For OmpA^Δ*CTD*^ chromosomal mutation, OmpA CDS of 190-346 was deleted with the reverse knocking-out primer was engineered to include a CDS of a his-6 tag and a stop codon. For Lpp^Δ*K58*^ mutation, the codon of K58 was deleted with forward primers included a stop codon after R57.

### Antibiotic susceptibility testing

Ampicillin minimum inhibitory concentrations (MIC) were determined for bacterial strains according to the Clinical and Laboratory Standards Institute guideline [52] using ampicillin MIC test strips (Liofilchem, 920031) according to manufacture’s protocol.

### Bacterial BS and vancomycin sensitivity assay

Bacterial survival spotting assay were performed as described previously [3]. Briefly, overnight bacterial cultures were adjusted to OD_600_ of 1 and 10-fold serial diluted to 10^-7^ with fresh LB media. A 4 μl of each dilution preparations were spotted onto LB agar plates with indicated supplements where appropriate.

### Bacterial growth kinetic assay

Bacterial growth kinetics were recorded as previously outlined [3]. In brief, overnight bacterial cultures were diluted 1:200 into 200 μl of fresh LB media, with indicated supplements where appropriate, in a 96-well plate. The plates were incubated at 37 °C with aeration in a CLARIOstar plate reader (BMG, Australia), which was set to measure the optical density at 600 nm (OD_600_) every 6 min for a duration of 18 h.

### LPS silver staining

LPS silver staining was conducted following previously established methods [3]. Briefly, bacterial cells (10^9^ CFU) harvested during mid-exponential growth were collected by centrifugation (20,000×*g* for 1 min) and lysed in 50 μl of SDS sample buffer, followed by heating at 100 °C for 10 minutes. The samples were then cooled to room temperature and treated with 50 μg/ml proteinase K (PK, NEB) for 18 h at 60 °C. After this treatment, the samples were heated again at 100 °C for 10 min, and 2-5 μl of each sample was loaded onto 10-20% SDS-tricine gels (Invitrogen, #EC66252BOX), with LPS being silver stained as previously described [53].

### Colicin E2 DNA endonuclease sensitivity assay

Determination of colicin E2 DNA endonuclease sensitivity was carried out as previously described [3]. Briefly, overnight bacterial culture was adjusted to OD_600_ of 0.5, and spread onto a LB agar plate using a sterile cotton tip applicator. Plates were allowed to dry at room temperature, and a 5 μl of purified colicin E2 [3] diluted in 2-fold series in PBS was spotted onto the plate. Plates were incubated overnight at 37°C for 18 h and the sensitivity level was determined by the minimum colicin E2 concentration that showed clear bacterial growth inhibition.

### MG1655-SΔwaaL BS suppressor mutant (BP) library acquisition and profiling

Suppressor mutants of MG1655-SΔ*waaL* resistant to DOC were obtained by spreading an overnight culture of MG1655-SΔ*waaL*, grown in LB (diluted 1:2,000 in fresh LB media), onto LB agar plates supplemented with 2.5 mM DOC. The plates were then incubated overnight at 37 °C. A total of 157 independent suppressor mutants (designated as BP library) were isolated and carefully streaked onto non-selective LB plates, which were confirmed with no observable growth defects and subsequently stored at -80 °C for further analysis.

To rank the DOC resistance levels of the MG1655-SΔ*waaL* suppressor mutants in BP library, overnight cultures of the bacterial strains, grown in LB media at 37 °C without any observable growth defects, were diluted 1:100 in 200 μl of LB media supplemented with 2.5 mM DOC in 96-well plates. These plates were incubated at 37 °C for 6 h and OD_600_ was recorded every 6 min as described above, and mutants were ranked according to OD_600_ readings at the 3.5-h mark, with higher OD_600_ values indicating greater resistance to DOC.

### Construction of waaL-complemented suppressor mutant (SBP) sub-library and selection

To exclude the suppressor mutants with mutations abolishing OAg biosynthesis and assembly, the primary suppressor library was complemented by pWaaL [3] via polyethyleneglycol (PEG) chemical transformation [54]. Briefly, MG1655-SΔ*waaL* suppressor mutants were grown to an OD_600_ of 0.2 in 1 ml LB media, were harvested via centrifugation (5,000×*g*, 4 °C). Bacterial cell pellets were mixed with 100 μl of ice-cold TSS buffer [LB medium, 10% (w/v) PEG (MW ∼3,500 Da), 10 mM MgCl_2_ and 10 mM MgSO_4_, pH 6.5] supplemented with 1 ng of pWaaL and incubated on ice for 30 min. After incubation, a 900 μl of LB supplemented with 0.2% glucose were added and the reaction was further incubated at 37 °C shaking for 1h before plating out on LB agar plates supplemented with appropriate antibiotic. Plates were incubated for 18 h at 37 °C to acquire transformants. All 158 MG1655-SΔ*waaL* suppressor mutants yielded transformants which were stored and designated as SBP library.

To identify mutants that potentially harbor suppressor mutations abolishing OAg biosynthesis and assembly, strains from SBP library were patched onto LB agar plates supplemented with 100 μg of ColE2, 2.5 mM DOC or LB agar plates pre-spread with 100 μl of P1kc phage lysate at 10^11^ pfu, respectively. Plates were incubated at 37 °C to observe sensitivity to ColE2, DOC and P1kc phage. Corresponding SBP mutants of top-ranked (top 31) BP suppressor mutants according to DOC resistance were further validated via LPS silver staining and ColE2 sensitivity assay detailed above. Suppressor mutants with identified defects in OAg biogenesis was excluded for whole genome sequencing analysis.

### Whole genome sequencing

For whole genome sequencing of bacterial strains, genomic DNA from MG1655-SΔ*waaL*, and suppressor mutants was extracted using the Qiagen QIAamp DNA Blood Mini Kit, following the manufacturer’s instructions. Samples were then prepared for DNBseq DNA library construction (BGI) and subsequently sequenced using DNBSEQ PE150 technology (BGI). The processed reads for each strain were mapped to the NCBI MG1655 reference genome (Accession number U00096) using Geneious 8.0. Sequence variations between our laboratory strain of MG1655 and the online reference genome determined and reported previously [3] were excluded.

### Western immunoblotting

For Western immunoblotting, whole bacterial cell lysate was prepared in SDS-sample buffer and heated at 95 °C for 5 min. Protein was separated via SDS-PAGE and transferred onto nitrocellulose membrane and detected with antibodies. Successful protein transfer was confirmed with ponceau S staining and imaged to serve as a loading control. For Western immunoblotting of O16 LPS, polysaccharide samples separated by SDS-tricine gel electrophoresis and processed as above. Rabbit polyclonal anti-O16 antibodies were purchased from SSI Diagnostica (#SSI85012), rabbit polyclonal anti-OmpA is a generous gift from Prof Ulf Henning (MaxPlank Institute for Biology Tübingen), rabbit polyclonal anti-SurA, rabbit polyclonal anti-Skp are generous gifts from Prof Thomas Silhavy (Princeton University), rabbit polyclonal anti-MBP-DegP is a generous gift from Prof Michael Ehrmann (University of Duisburg-Essen), rabbit anti-AcrB is gifted by Prof Reiter Venter (University of South Australia). Anti-Lpp antibody was affinity purified from anti-OmpF/C/A antibodies that are generous gifts from Prof Rajeev Misra (Arizona State University). Briefly, anti-OmpF/C/A antibody (1/100 in PBS) was incubated with nitrocellulose membrane area transferred with Lpp protein from whole cell lysate overnight at 4 °C. Membranes were washed three times with PBS containing 0.5% (v/v) tween 20, and eluted with 100 μl of 0.1 M glycine, pH 3.0. Eluted antibodies were pH adjusted to pH 7.0 and verified with Western immunoblotting.

### Protein sequence alignment and structure comparison analysis

The BamA structure (PBD 7TTC) [55] and LptD structure (PDB 4Q35) [56] were analysed and annotated by using UCSF Chimera X [57].

### Membrane fractionation by sucrose density gradient centrifugation

Bacterial cell membrane fractionation was performed as described previously [58] with modifications. Briefly, overnight bacterial culture was subculture 1:100 in 100 ml LB and grown for 2 h at 37 °C until OD_600_ reached 0.8. DOC was added to 0.5 mM and allowed for another 1 h growth. Cells were harvested via centrifugation at 5,000 g, and the resulting cell pellet was resuspended in 5 ml osmotic buffer [0.5 M sucrose, 10 mM Tris pH 7.5], treated with lysozyme (200 μg/ml) and 1 mM EDTA on ice for 10 min and sonicated to lyse cells (with approx. 1,000 J accumulative energy). Sonicates were briefly centrifuged at 5,000×*g*, 4 °C and the resulting supernatant was ultracentrifuge at 140,000◊*g* (Beckman Coulter type 70.1 rotor with tube #355603) for 1.5 h. The resulting membrane pellet was then dissolved in 1 ml Low-density isopycnic sucrose gradient solution [20% w/v sucrose, 1 mM EDTA, 1 mM Tris pH 7.5]. Centrifuge tube was filled with 2 ml of [73% w/v sucrose, 1 mM EDTA, 1 mM Tris pH 7.5], then 4 ml of [45% w/v sucrose, 1 mM EDTA, 1 mM Tris pH 7.5], then 1 ml of solubilized membrane fraction, and finally topped up with ∼6 mL of [20% w/v sucrose, 1 mM EDTA, 1 mM Tris pH 7.5] until tubes is full. Centrifugation was performed at 100,000 ◊*g* (Beckman Coulter SW40 Ti) for 17 h with slow acceleration and no brake at 4 degrees. Tubes with separated membranes were imaged. Fractions (1 ml) were collected by 1 ml P1000 tip (with tip cut by ∼5 mm). For NADH dehydrogenase assay, each isolated fractions were diluted 1/10 in 10 mM Tris pH7 buffer (10 μl+90 μl) in a 96-well plate, followed by the addition of 10 μl of 10 mg/ml NADH (in MQ water), and plates were immediately read at 340 nm every 30 s for 40 min. Readings at 20 min were compared with low optical density at 340 nm indicating high enzyme activity of NADH dehydrogenase. All fractions were subjected to Western immunoblotting as described above and detected with anti-OmpA, anti-AcrB and anti-Lpp antibodies.

### Peptidoglycan purification and quantification of muropeptides

Peptidoglycan isolation and purification was described previously [59]. Briefly, bacterial cell culture (400 ml) grown to OD_600_ of 0.8 were supplemented with 2.5 mM DOC for 30 min, cooled rapidly on ice-water bath followed by centrifugation (10,000◊*g*, 10 min, 4 °C). Cell pellet was resuspended in 6 ml of ice-cold water and dropped into boiling 8% (w/v) SDS solution. The samples continued to be boiled for 30 min before subjected to ultracentrifugation (437,000◊*g*, 1 h) with repeated washes with water to remove SDS. The SDS-free pellet was resuspended in 10 mM Tris-HCl (pH 7.0), 320 mM imidazole and incubated with 150 μg/ml α-amylase (Sigma) at 37 °C for 2 h, followed by incubation with 200 μg/ml pre-heated pronase E (Sigma) at 60 °C for 2 h. The sample was then mixed with an equal part of 4% SDS and boiled for 30 min. PG was then harvested via ultracentrifugation followed by washes to remove SDS as described above. Purified PG was digested with 50 μg/ml Cellosyl (gift from Hoechst, Frankfurt, Germany) in 20 mM sodium phosphate (pH 4.8) overnight at 37 °C, followed by heating at 100 °C for 10 min. Soluble muropeptides were harvested via brief centrifugation (20,000 ◊*g*, 1 min) and mixed with an equal volume of 0.5 M sodium borate (pH 9.0) and reduced with sodium borohydride (NaBH_4_ ) for 30 min at room temperature. The pH was adjusted to 3.5-4.5 with 20% phosphoric acid. The samples were stored at −20°C.

Separation of the reduced muropeptides by HPLC was described previously [59]. The eluted muropeptides were monitored by measuring absorbance at 204 nm. Relative quantification of each muropeptide species was performed by integration of the peaks of the HPLC profile, and the muropeptides were grouped into classes according to structural similarities [59]. Chromatograms are included in supplementary materials (S7 Fig).

## Acknowledgement

This work is primarily funded by an Early Career Research Ideas Grant from the Faculty of Health, Queensland University of Technology (Australia) to JQ, and in part by an Australian Research Council project grant (DP210101317), the Max Planck Queensland Centre on the Materials Science of Extracellular Matrices to MT, and start-up funds from the University of Queensland to WV. The Ian Potter Foundation sponsored the CLARIOStar high-performance microplate reader (BMG, Australia) and epifluorescence microscope. The funders had no role in study design, data collection and analysis, decision to publish, or preparation of the manuscript. We thank Dr Daniela Vollmer (University of Queensland) for providing technical trainings in analysing the peptidoglycan composition.

## Author contributions

JQ conceptualised the project; JQ contributed to experimental design; JQ conducted all experiments, and contributed to data collection and analysis; JQ, WV and RM contributed to data interpretation; JQ and MT obtained the funding. YH contributed to the generation of experimental materials. JQ wrote the manuscript, and all authors edited the manuscript.

## Data availability

All data generated or analysed during this study were included in this article and supplementary files.

---

**S1 Fig.**
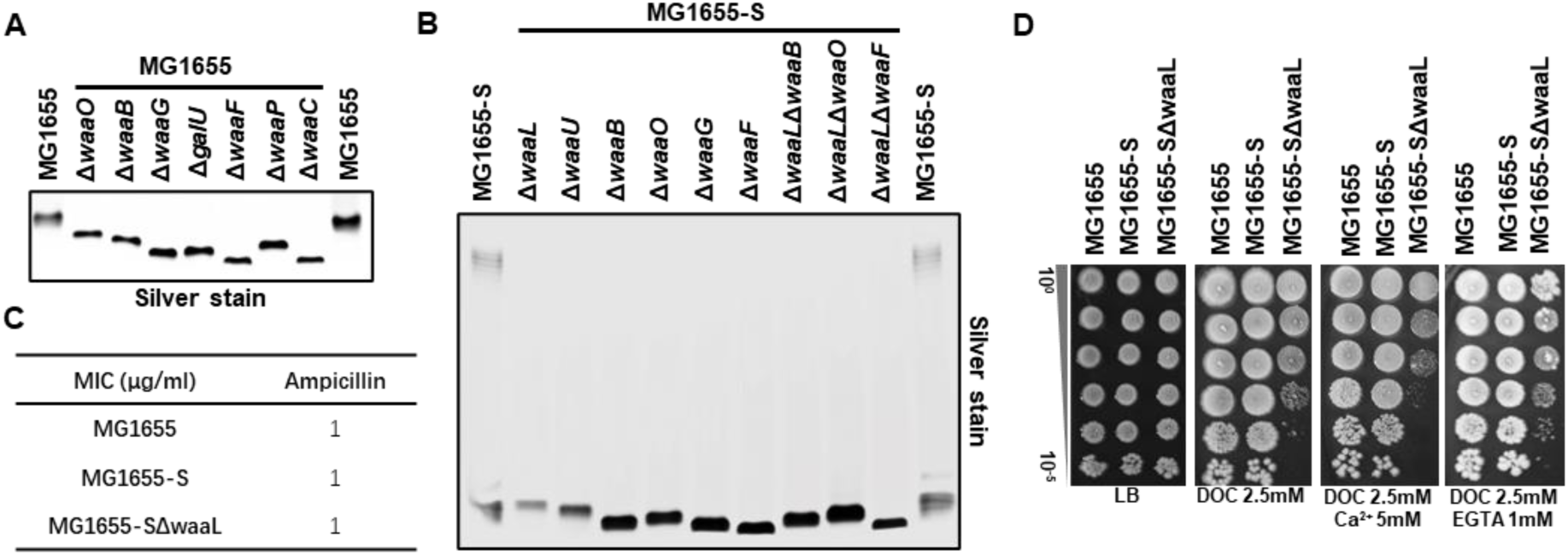
Validation of LPS core truncations mutants of MG1655 derivative strains and effect of LPS-Ca^2+^ bridge in BS resistance. **(A&B)** LPS silver staining of proteinase K-treated whole cell lysates of indicated MG1655 derivative strains. (**C**) MIC of ampicillin determined for indicated *E. coli* K-12 strains. (**D**) BS sensitivity assay of indicated *E. coli* K-12 strains in the presence of 5 mM Ca^2+^ (CaCl_2_) or 1 mM EGTA.

**S2 Fig.**
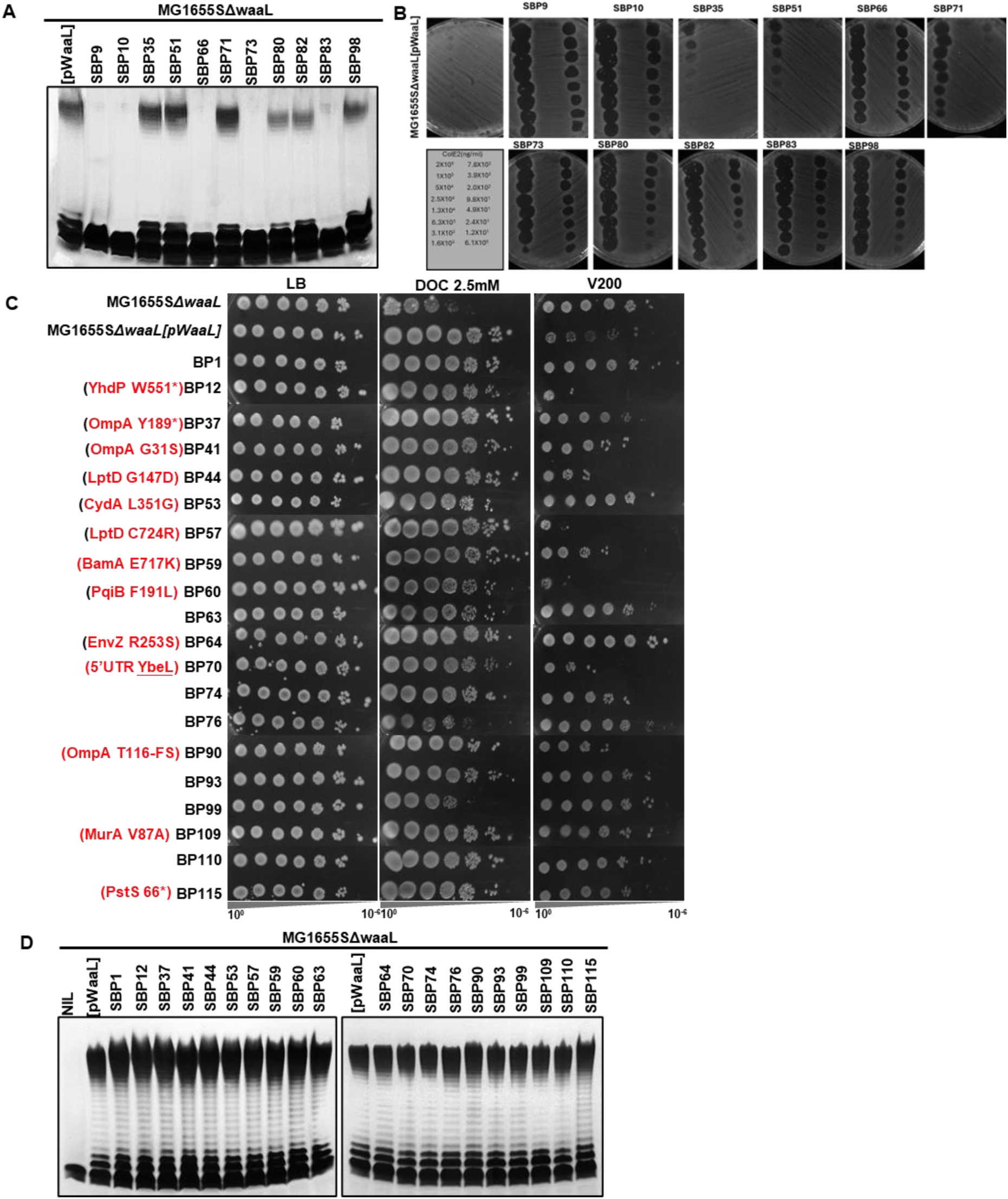
Characterisation of LPS profile of SBP mutants and their corresponding BP suppressor mutants’ BS resistance. (**A**) LPS silver staining of proteinase K-treated whole cell lysates of indicated MG1655-SΔ*waaL* SBP mutant strains. (**B**) Colicin E2 sensitivity assay of indicated MG1655-SΔ*waaL* SBP mutant strains. Colicin E2 (5 μl, at indicated concentrations) was spotted onto LBA plate pre-spread with cultures of indicated MG1655-SΔ*waaL* SBP mutant strains. Sensitivity to Colicin E2 was shown as inhibited bacterial growth. (**C**) BS and vancomycin (200 μg/ml) sensitivity assay to indicated MG1655-SΔ*waaL* BP mutant strains. Mutations mapped through WGS was indicated in red. (**D**) LPS silver staining of proteinase K-treated whole cell lysates of indicated MG1655-SΔ*waaL* SBP mutant strains.

**S3 Fig.**
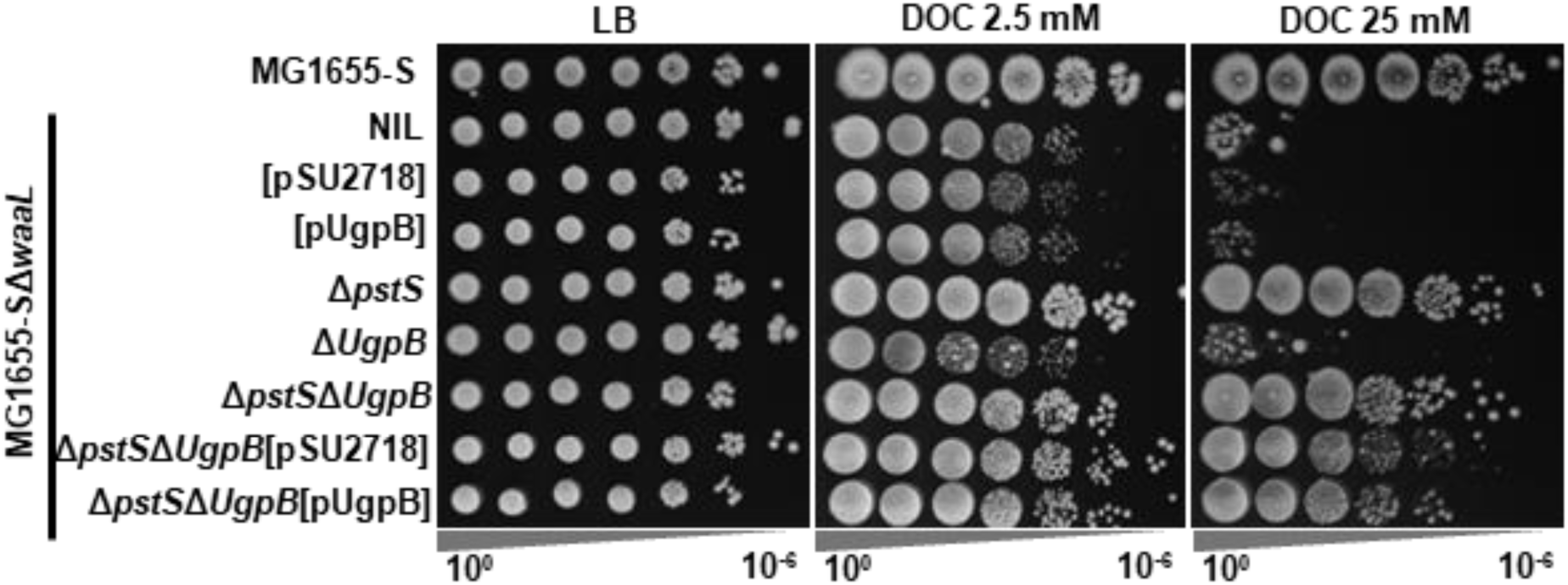
Investigation the role of UgpB in conferring BS resistance to MG1655-SΔ*waaL.* Indicated MG1655-S derivative strains harbouring indicated expression construct were assayed for BS resistance with 2.5 mM and 25 mM DOC.

**S4 Fig.**
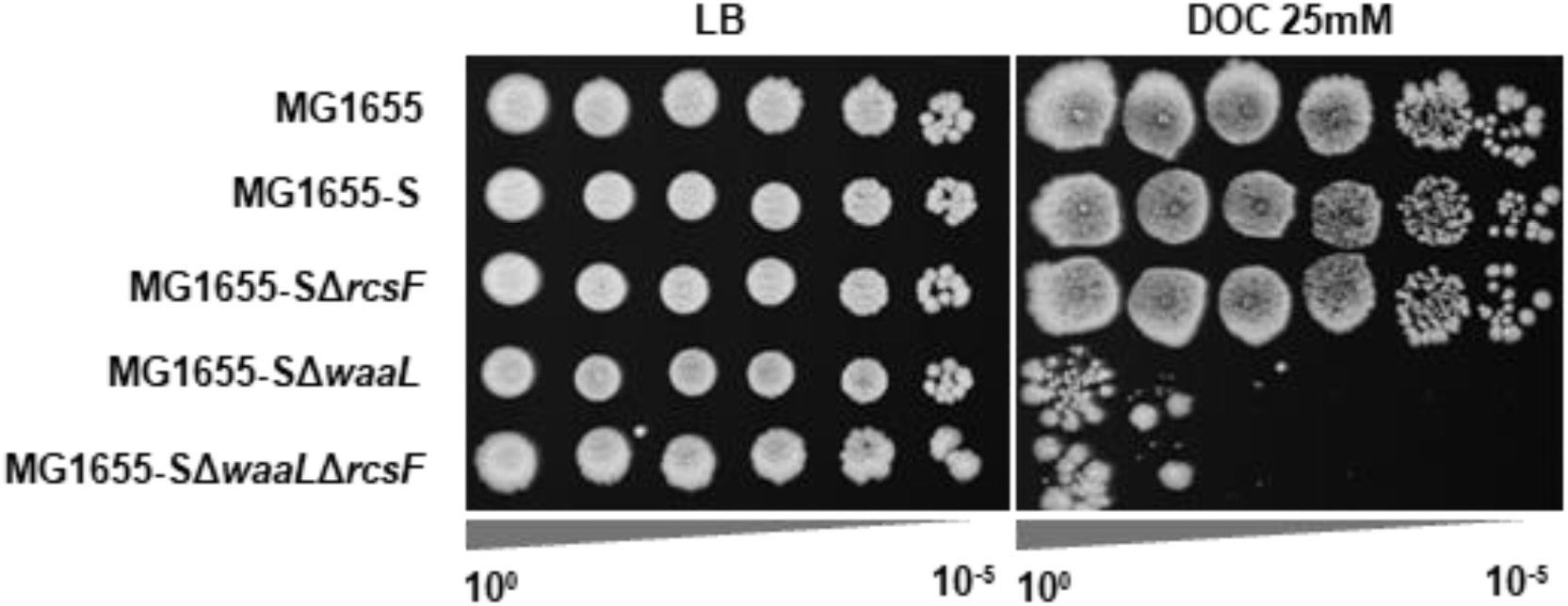
Investigation the role of RcsF in conferring BS resistance to MG1655-SΔ*waaL.* Indicated MG1655-S derivative strains were assayed for BS resistance with 25 mM DOC.

**S5 Fig.**
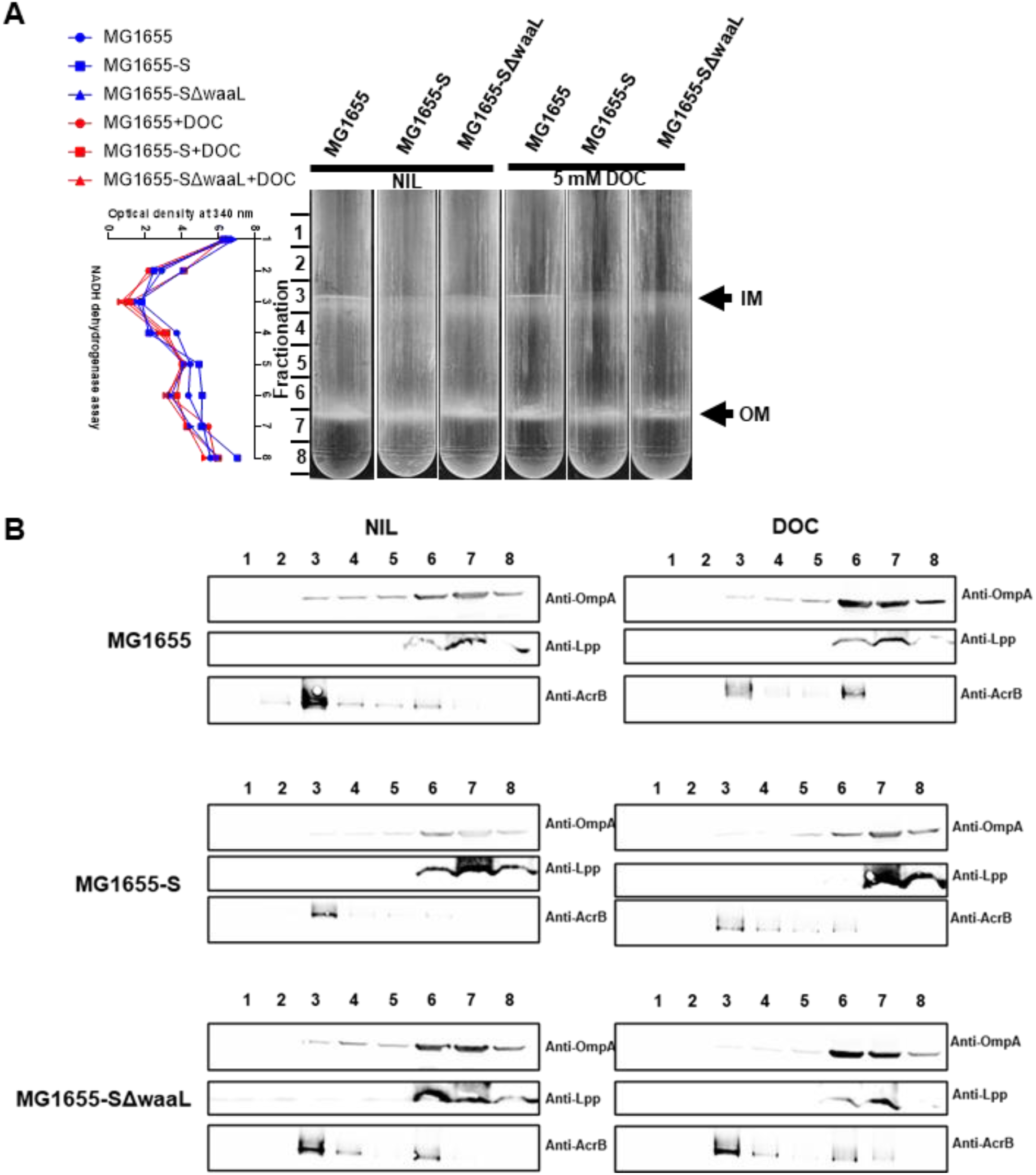
Membrane localisation of Lpp in MG1655-SΔ*waaL* grown in the presence of BS. Bacterial cell membranes were isolated from indicated bacterial cell cultures with or without 5 mM DOC (detailed in Materials and Methods). IM and OM (marked in **A**) were separated via sucrose gradient density centrifugation (**A**). The separated membrane gradient was fractionated (1 ml) indicated on the left side. Samples of fractions were taken to evaluate the NADPH dehydrogenase activity to validate the successful separation of IM from OM, with lower OD_340_ reading correlates to higher activity of NADPH dehydrogenase detected in the fraction (**A left curve**). Validated fractions were then subjected to Western immunoblotting (**B**) to investigate the membrane localisation of Lpp via anti-Lpp antibodies. Anti-OmpA and anti-AcrB were used as a reference for OM and IM localisations, respectively.

**S6 Fig.**
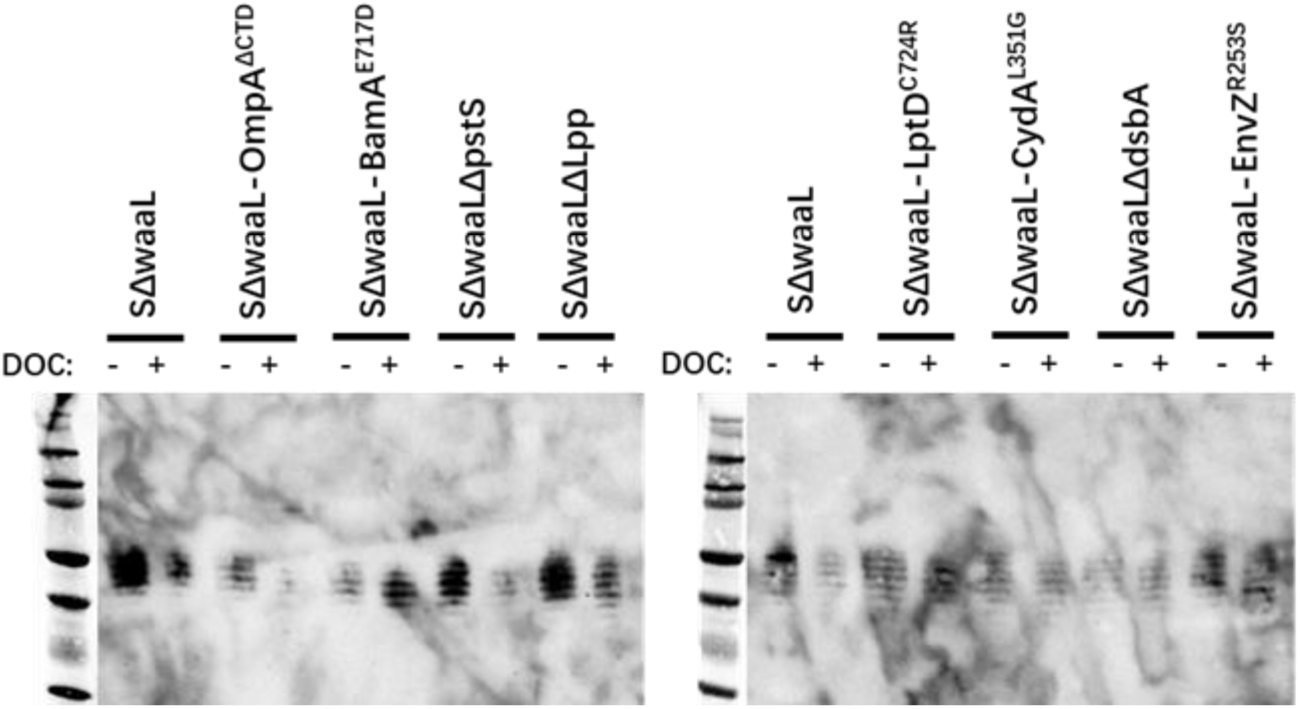
Confirmation of presence of UndPP-OAg in the MG1655-SΔ*waaL* derivative strains via Western immunoblotting. MG1655-SΔ*waaL* SBP mutant strains were allowed to grow to OD_600_ of 0.8, and DOC (25 mM) was added for another 30 min growth. MG1655-SΔ*waaL* showed clear lysis, and all suppressor strains showed mild lysis. Proteinase K-treated bacterial cell lysates were prepared and then subjected to Western immunoblotting with anti-O16 antibodies. UndPP-OAg was detected as ladder pattern bands.

**S7 Fig.**
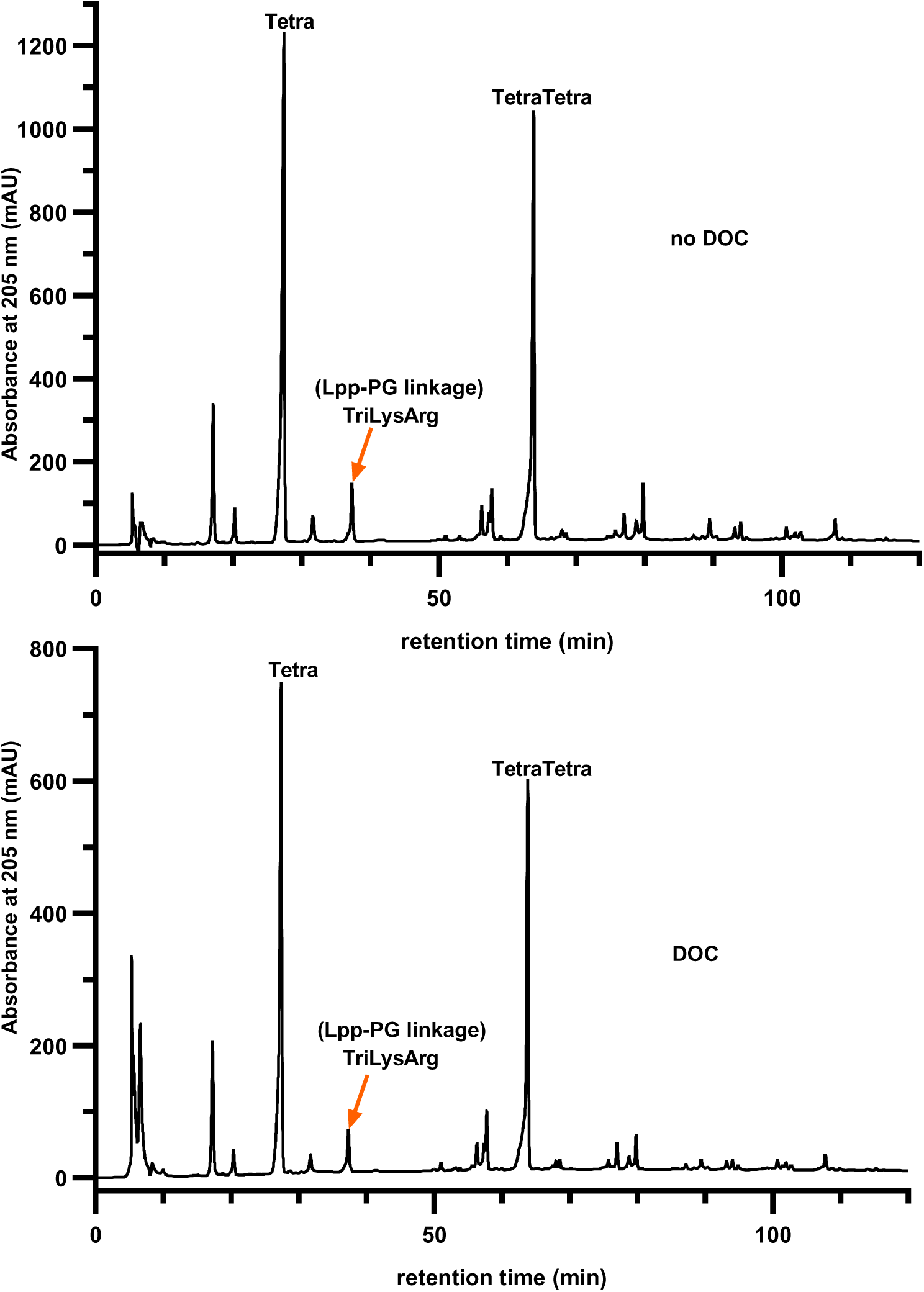
Muropeptide profile of MG1655-SΔ*waaL* grown in the presence or absence of BS. Chromatography of muropeptide derived from MG1655-SΔ*waaL* grown in the absence (**A**) or presence (**B**) of DOC were isolated and processed according to Materials and Methods, the processed muropeptides were profiled by HPLC as detailed in Materials and Methods. Major peaks corresponding to Tetra, Tetra-Tetra and TriLysArg (LPP-PG linkage) are indicated according to Glauner (59).

**Supplementary Table 1.** BS resistance rank of MG1655-SΔwaaL BP suppressor mutants and characterization of its derivative SBP mutants in ColE2 and P1 resistance.

**Supplementary Table 2.**
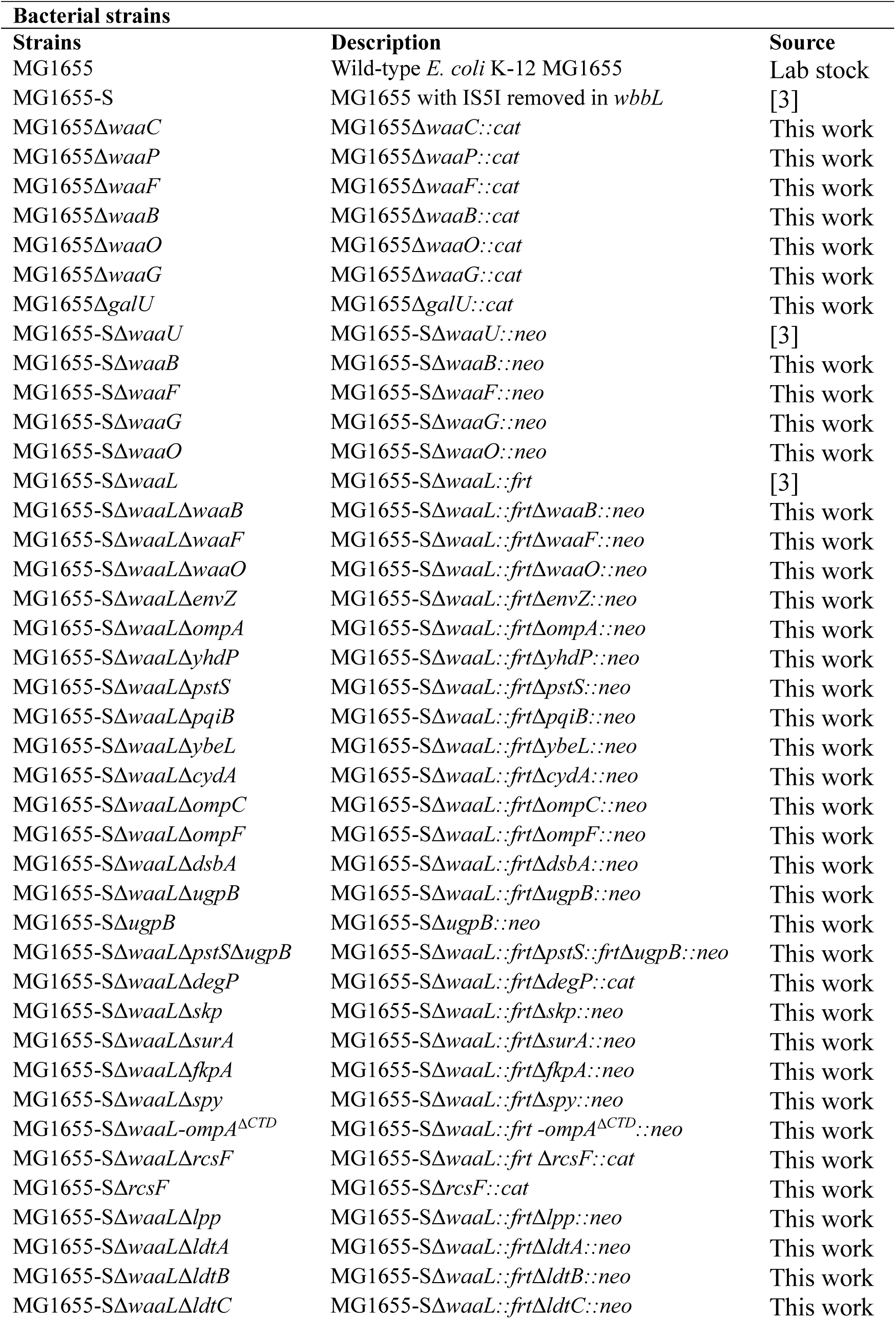
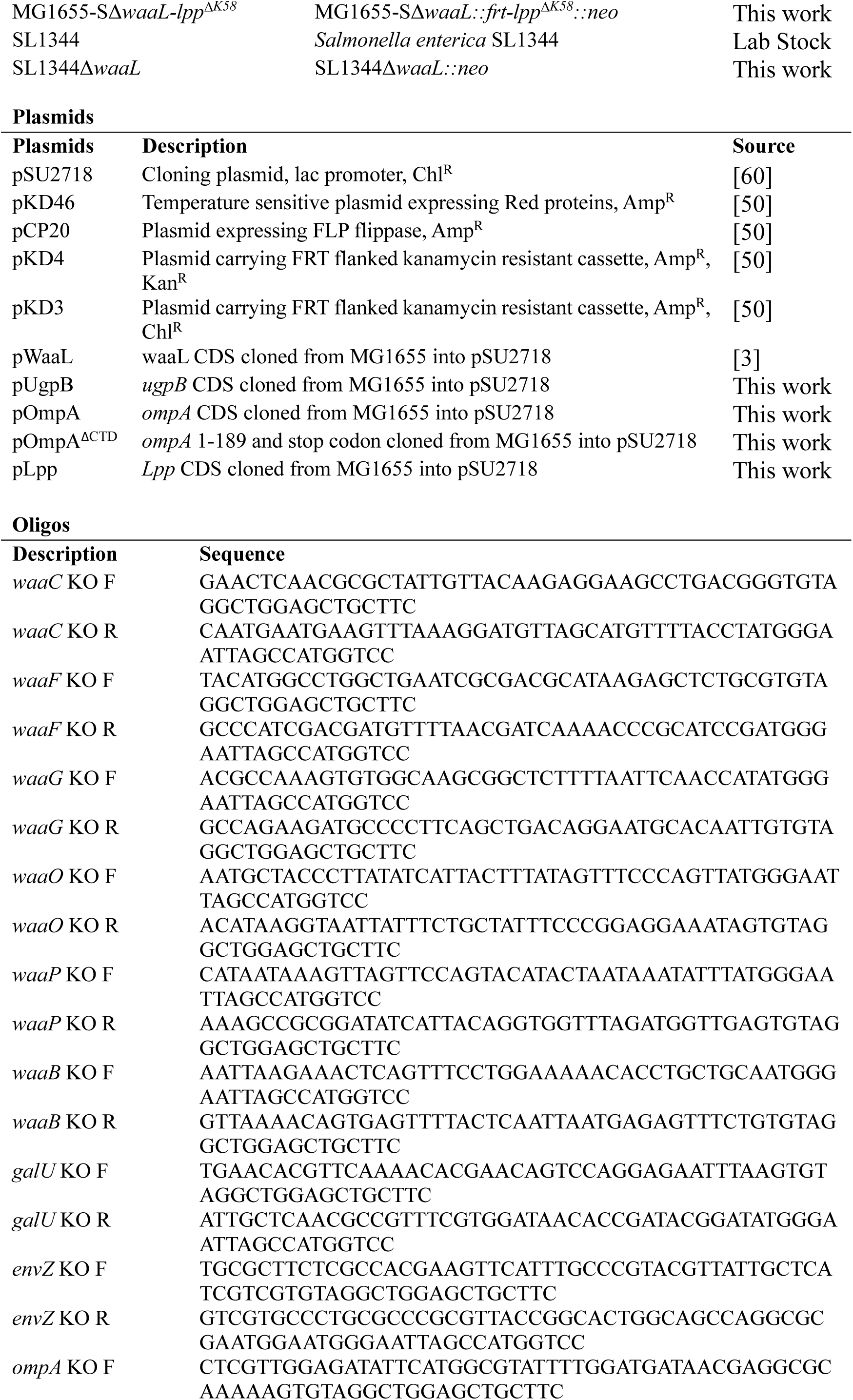
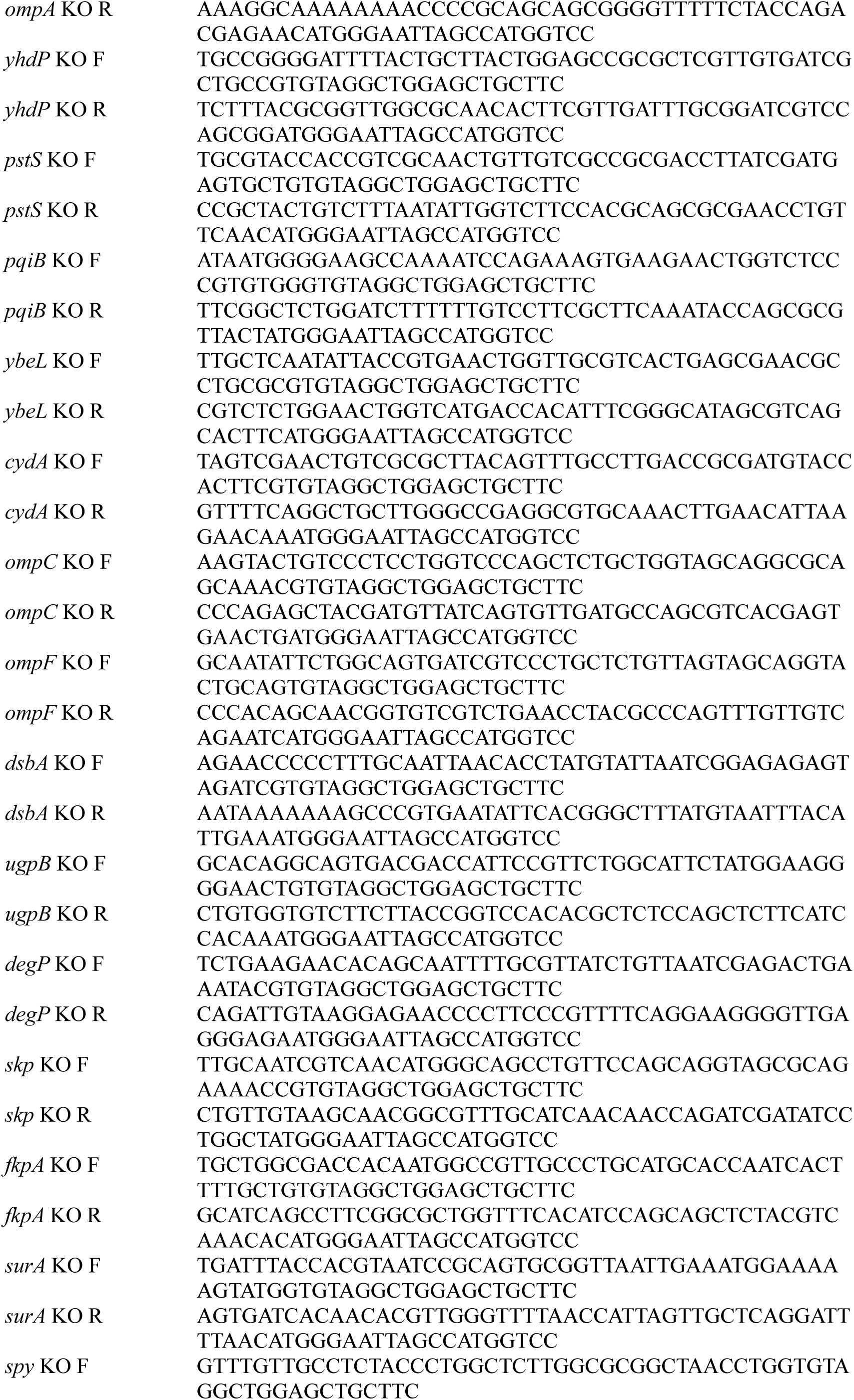
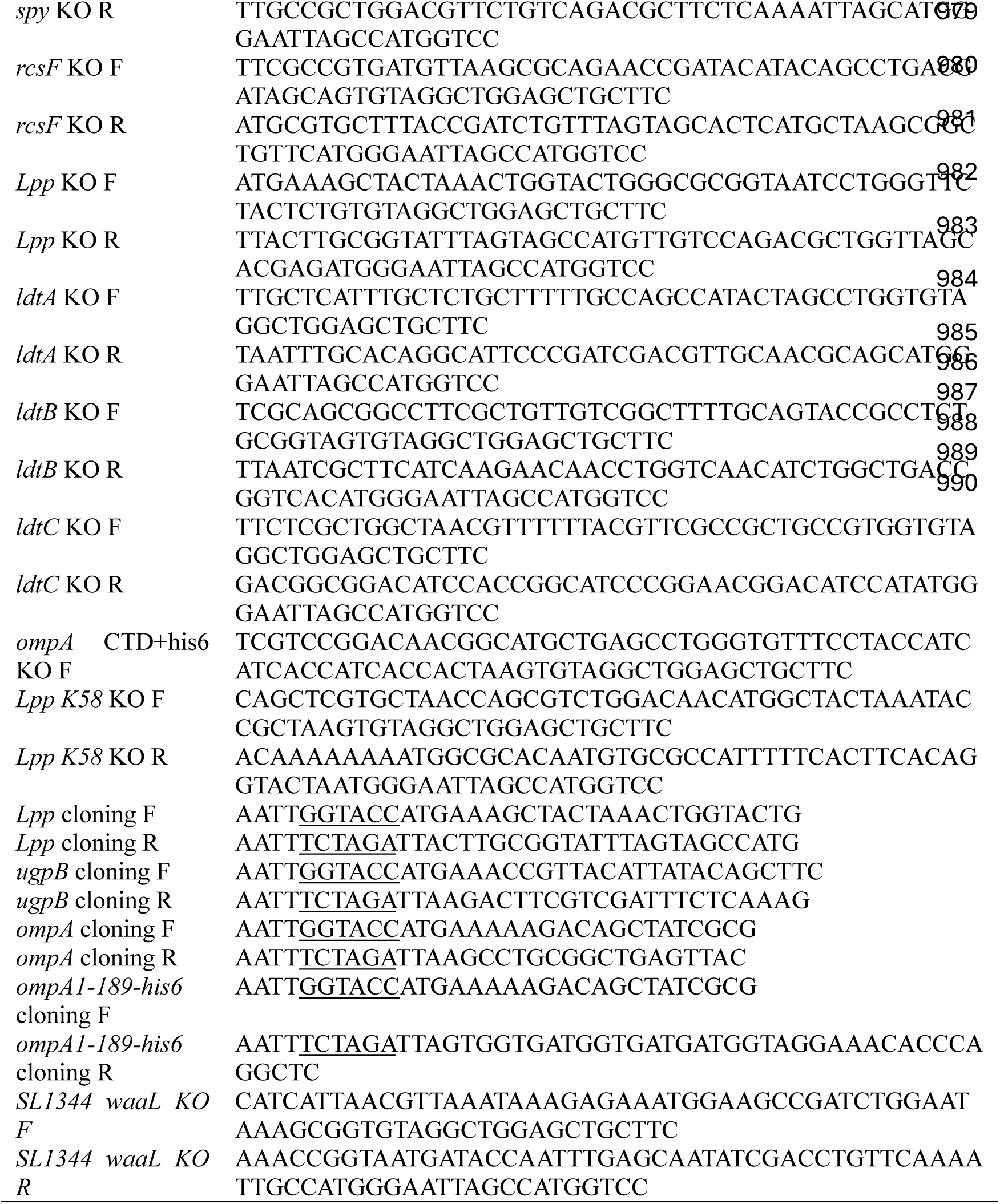
Strains, plasmids, and oligonucleotides.

**Supplementary Table 3.** Summary of muropeptide profile of MG1655-SΔ*waaL* grown in the absence or presence of BS.

